# Transient Impairment in Microglial Function Causes Sex-Specific Deficits in Synaptic and Hippocampal Function in Mice Exposed to Early Adversity

**DOI:** 10.1101/2024.02.14.580284

**Authors:** Sahabuddin Ahmed, Baruh Polis, Sumit Jamwal, Basavaraju G. Sanganahalli, Zoe MacDowell Kaswan, Rafiad Islam, Dana Kim, Christian Bowers, Lauryn Giuliano, Thomas Biederer, Fahmeed Hyder, Arie Kaffman

**Affiliations:** Department of Psychiatry, Yale University School of Medicine, 300 George Street, Suite 901, New Haven CT, 06511, USA; Department of Radiology & Biomedical Imaging and Magnetic Resonance Research Center, Yale University, New Haven, CT, 06520, USA; Department of Biomedical Engineering, Yale University, New Haven, CT, 06519, USA; Department of Neurology, Yale School of Medicine, 100 College Street, New Haven, CT 06510, USA

**Keywords:** Early life adversity, mice, microglia, astrocytes, synaptic pruning, limited bedding and nesting, hippocampus

## Abstract

Abnormal development and function of the hippocampus are two of the most consistent findings in humans and rodents exposed to early life adversity, with males often being more affected than females. Using the limited bedding (LB) paradigm as a rodent model of early life adversity, we found that male adolescent mice that had been exposed to LB exhibit significant deficits in contextual fear conditioning and synaptic connectivity in the hippocampus, which are not observed in females. This is linked to altered developmental refinement of connectivity, with LB severely impairing microglial-mediated synaptic pruning in the hippocampus of male and female pups on postnatal day 17 (P17), but not in adolescent P33 mice when levels of synaptic engulfment by microglia are substantially lower. Since the hippocampus undergoes intense synaptic pruning during the second and third weeks of life, we investigated whether microglia are required for the synaptic and behavioral aberrations observed in adolescent LB mice. Indeed, transient ablation of microglia from P13-21, in normally developing mice caused sex-specific behavioral and synaptic abnormalities similar to those observed in adolescent LB mice. Furthermore, chemogenetic activation of microglia during the same period reversed the microglial-mediated phagocytic deficits at P17 and restored normal contextual fear conditioning and synaptic connectivity in adolescent LB male mice. Our data support an additional contribution of astrocytes in the sex-specific effects of LB, with increased expression of the membrane receptor MEGF10 and enhanced synaptic engulfment in hippocampal astrocytes of 17-day-old LB females, but not in LB male littermates. This finding suggests a potential compensatory mechanism that may explain the relative resilience of LB females. Collectively, these studies highlight a novel role for glial cells in mediating sex-specific hippocampal deficits in a mouse model of early-life adversity.

## INTRODUCTION

Adverse childhood experiences, such as abuse, neglect, extreme poverty, and neighborhood violence, can lead to abnormal brain development and early-life emotional and cognitive challenges (Kaffman and Meaney, 2007; McLaughlin et al., 2020; Teicher and Samson, 2016). Approximately half of all childhood psychopathologies are attributed to early life adversity (ELA) (Green et al., 2010). In many cases, these early consequences persist into adulthood, resulting in chronic psychiatric and medical conditions that are difficult to treat (Anda et al., 2006; Kaffman and Meaney, 2007; Nemeroff, 2016). A better understanding of the underlying biology is essential for diagnosing the structural and functional changes associated with ELA and for developing more effective interventions.

Some of the most consistent findings in individuals exposed to ELA are reduced hippocampal volume and abnormal hippocampal function (Crozier et al., 2014; De Bellis et al., 2013; Lambert et al., 2019; Teicher and Samson, 2016), with some evidence indicating more pronounced deficits in men than women (Garvin and Bolton, 2022; Teicher and Samson, 2016; White and Kaffman, 2019). Rodents exposed to different models of ELA also exhibit abnormal hippocampal function, with particularly pronounced deficits observed in mice reared with limited bedding and nesting (LB) (Rocha et al., 2021). However, most of the studies to date have focused on outcomes in adult LB males, with only a few examples examining this issue in both sexes (Bath et al., 2017; Naninck et al., 2015; Rocha et al., 2021). Additionally, only two studies have examined the effect of LB on hippocampal function and development in adolescent mice (Bath et al., 2017; Islam et al., 2023), a period in which sex-specific effects of ELA appear to be more prominent (Gershon et al., 2008; White and Kaffman, 2019).

Microglia, the immune innate cells in the brain, control the refinement of connectivity and play a critical role in establishing sexually dimorphic processes during brain development (VanRyzin et al., 2020). Microglia are also responsible for distinct physiological and behavioral responses in adult males and females (Dorfman et al., 2017; Sorge et al., 2015; Villa et al., 2018; Yanguas-Casás, 2020). Given the prominent role that microglia play in programing sex differences we investigated whether abnormal microglial-mediated synaptic pruning during the second and third weeks of life contributes to sex-specific deficits in synaptic function and contextual freezing observed in adolescent LB mice. Microglia play a crucial role in eliminating nonfunctional synapses during a critical period of development and disrupting this process leads to the retention of a large number of weak synapses, reduced connectivity, and abnormal behavior and cognition later in life (Bolton et al., 2022; Filipello et al., 2018; Johnson and Kaffman, 2017; Zhan et al., 2014). Moreover, significant impairments in microglial-mediated synaptic pruning occur in LB pups, as well as increased spine density that persists in adolescent mice (Bolton et al., 2022; Dayananda et al., 2022). In this study, we determined that LB causes severe synaptic and contextual fear conditioning deficits in adolescent male, but not female, littermates. These deficits were accompanied by impairment in microglial-mediated phagocytic activity in 17-day-old male and female pups when synaptic pruning peaks in the developing hippocampus (Filipello et al., 2018; Scott-Hewitt et al., 2020; Zhan et al., 2014). Abnormal microglial phagocytic activity did not persist into adolescence when phagocytic activity is eight-fold lower. Furthermore, transient elimination of microglia during the second and third weeks of life induced similar sex-specific deficits in synaptic function and contextual fear conditioning observed in LB mice and chemogenetic activation of microglia during this critical period was sufficient to normalize the synaptic and contextual deficits observed in adolescent LB male mice. Finally, we found that LB female, but not LB male littermates are able to upregulate synaptic pruning in astrocytes.

## METHODS

### Animals

BALB/cByj mice (Stock # 001026, Jackson Laboratories) were housed in standard Plexiglas cages and kept on a standard 12:12 h light-dark cycle (lights on at 07.00 AM) with constant temperature and humidity (22 °C and 50%) and food and water provided ad libitum CX3CR1^creET2^-DTA mice (C57BL/6J background) were generated by crossing CX3CR1-Cre-ET2 mice (Jax 021160) and ROSA26-eGFP-DTA mice (Jax# 006331). CX3CR1 ^creET2^-Gq-DREADD mice (C57BL/6J background) were generated by crossing CX3CR1-Cre-ET2 mice (Jax 021160) and CAG-LSL-Gq-DREADD mice (Jax# 026220). All studies were approved by the Institutional Animal Care and Use Committee (IACUC) of Yale University and were conducted in accordance with the recommendations of the NIH Guide for the Care and the Use of Laboratory Animals.

### Limited Bedding

The limited bedding (LB) procedure was performed as described previously (Dayananda et al., 2022; Johnson et al., 2018; White et al., 2020). Briefly, mice were mated at a 3:1 female to male ratio, in standard mouse Plexiglas cages layered with 500 cc of corncob bedding but with no nesting material. Visibly pregnant dams were transferred to maternity cages containing 500 cc corncob bedding but no nesting material. At birth, postnatal day (P0) litters were culled to 5-8 pups and randomized to either control (CTL) or LB conditions. Mice raised under CTL condition were provided with 500 cc of fresh corncob bedding with an additional 15 cc of soiled bedding from the original cage and one 5 × 5 cm nestlet per cage. LB litters were provided with 125 cc of corncob,15 cc of soiled bedding from the original cage, and no nesting material. Bedding was changed weekly, and mice were weaned on P26 and housed with 2-3 individuals of the same-sex and condition per cage.

### Contextual fear conditioning

Contextual fear conditioning was performed using a Med Associates’ fear conditioning chamber as previously described (White et al., 2020).

### Tissue collection

Tissues were collected between 11:00 and 13:00 to minimize the diurnal effects of corticosterone. Mice were anesthetized and transcardially perfused with ice-cold PBS/heparin (50 u/ml) solution (Bio-Rad, Cat #1610780; Sigma, Cat# H3393), followed by 10% formalin (Polyscience, Cat# 08279-20). The brains were postfixed for 1hr with 10% formalin at r.t. and then stored in PBS at 4°C until they were processed for immunohistochemistry or DiOlistic labeling.

### DiOlistic labeling

This procedure was performed as described previously (Forlano and Woolley, 2010) with the following modifications. Perfused brains were coronally sectioned at 200 µm using a VT1000S vibratome (Leica). Sections containing the dorsal hippocampus (Paxinos, Bregma -1.82 to - 2.30) were labeled with a Helios gene gun (Bio-Rad) using 200 psi and µm tungsten particles coated with fluorescent 1,1’-Dioctadecyl-3,3,3’,3’-Tetramethylindocarbocyanine Perchlorate (DiI, Thermo Fisher Scientific; D-282). The DiI was allowed to diffuse at r.t. for 24 hrs and then washed with PBS. The slices were then postfixed with 10% formalin for 2hrs at r.t, washed with PBS, mounted on Superfrost Plus slides (Therrmo Scientific, Cat. # 4951F-001) with Fluoromount-G^TM^ solution (Invitrogen, Cat. # 00-4958-02), and coverslipped.

### Immunohistochemistry

Fifty-micron coronal sections were collected using a VT1000S vibratome (Leica) in 6 pools, each containing 16-18 slices, spanning the entire rostral-caudal axis of the hippocampus. To assess microglial phagocytic activity, one pool of slices was stained with rabbit anti-Iba1 (1:500; Wako, Cat. #019-19741), mouse anti -PSD95 (1:100; Merck-Millipore, Cat. #MAB1596), and rat anti -CD68 (1:400; Bio-Rad, Cat. # MCA1957T).To assess the density of functional glutamatergic synapses, slice were incubated with guinea pig anti-Vglut2 (1:700; EMD-Millipore Cat. #AB2251-I), and mouse anti-PSD95 (1:100; Merck-Millipore, Cat. #MAB1596). Astrocytic phagocytic activity and MEGF10 levels were characterized using sequential staining in order to minimize cross reactivity between mouse and rat antibodies. Briefly, sections were washed with TBST (3 X 15 min each) and blocked with Triton-X 100 (0.5%) + normal goat serum (NGS) (5%) for 2 hr. Sections were then first incubated with mouse anti-PSD95 antibodies (1:100; Merck-Millipore, Cat. #MAB1596) overnight at 4°C, washed and incubated with Alexa Flour^TM^ 555 goat anti-mouse secondary antibody (1:400; Invitrogen Cat. # A21422). The sections were again re-blocked with 5% NGS for 2 hr (second serum blocking) and subsequently stained with rat anti-GFAP (1:500; Invitrogen Cat. #13-0300), and rabbit anti-Megf10 (1: 400; EMD-Millipore Cat. #ABC-10). Stained slices were then labelled with the appropriate fluorescently labeled secondary antibodies (1:400; Thermo Fisher) and then mounted on glass slides with VECTASHIELD HardSet antifade mounting medium with DAPI (Vector laboratories Cat# 10955).

### Microscopy and image analysis

Spine density and morphology were assessed in fully impregnated CA1 pyramidal cells located in the dorsal hippocampus and imaged with a Zeiss LSM 880 confocal workstation equipped with Airyscan using 20X objective at 0.5 µm intervals. Secondary and tertiary apical dendrites located in the stratum radiatum and were 20 ±3 µm in length and 1.0 ±0.2 μm in diameter were then cropped and modeled using the semiautomated filament tracer tool in Imaris 10.0 (Oxford Instruments). The default setting of the Classify Spines X Tension tool in Imaris was subsequently used to classify spines as stubby, mushroom, long thin, or filopodia and to calculate total spine density and maturity index= density of mushroom/all other spine categories. The data from 6 dendrites were averaged to obtain the total spine density and maturity index for each individual animal. Microglial volume, CD68 volume, and the number of PSD95 puncta inside microglia located in the stratum radiatum were determined as described previously (Dayananda et al., 2022). To assess glutamatergic spine density, high-resolution (1024 x 1024), 6-8 confocal Z stack images of the stratum radiatum were acquired using an Olympus FV-3000 microscope with a 60X objective, 2x digital zoom and 0.30 μm intervals for a total thickness of 15-20 μm. The acquired images were deconvoluted and processed using the Imaris version-9.9.1 (Oxford Instruments) according to the following protocol. A 25 μm x 50 μm x10 μm region of interest was selected using the cropped 3D function with each channel adjusted using gaussian filter, automated background subtraction, and gamma correction. A 3D reconstruction spots were created for each channel using the ‘spot function’ with an XY diameter of 0.2 μm and Z-axis elongation at 10 μm. Spots were selected based on the ‘Quality’ filter type with the center point and radius size set at 10. Finally, the spots which were in close proximity (0-250 μm apart) were considered functional synapses and were calculated using the ‘shortest distance to spot-spot’ filter. The densities obtained from 6-8 pictures were averaged to determine VGlut2, PSD95, and glutamatergic synapse densities for each mouse. Z-stack images of GFAP-positive cells localized in the stratum radiatum were acquired using an Olympus FV-3000 using a 60x objective, 2x digital zoom, and 0.30 μm intervals for a total thickness of 20 μm and processed using the 3D surface rendering function in Imaris. A 50 μm x 50 μm x 10 μm region of interest was copped using the 3D function and each channel adjusted via a Gaussian filter using automated background subtraction and gamma correction. GFAP and Megf10 surfaces were created using a semiautomated thresholds of 0.3 μm and 0.2 μm respectively. PSD95 puncta were detected and quantified using the ‘spot function’ with a threshold of 0.2 μm. Finally, the number of PSD95 puncta and Megf10 volume engulfed inside GFAP-positive cells were quantified using the ‘spot closed to surfaces’ function with a threshold set at zero. Measurements obtained from 5-6 astrocytes were averaged to determine cell volume, number of PSD95 puncta per cell, and volume of MEGF10 staining per cell for each mouse.

### Resting state fMRI (rsfMRI)

We used functional connectivity density (FCD) mapping to assess the local functional connectivity. FCD is a voxel-and degree-based metric, which identifies the number of correlated voxels to a base voxel without identifying the precise location of the correlated voxels. These metrics are degree based, as they are based on the number of voxels that a given voxel is strongly correlated with, or functionally ‘‘connected to.’’ Global FCD determination is a standard graph theory-based analysis determining brain functional connectivity using resting state fMRI and originally demonstrated in the human brain (Tomasi and Volkow, 2010). Local functional connectivity was determined as described previously (Farina et al., 2021; Sanganahalli et al., 2021). Mice were initially anesthetized with 2-3 % isoflurane, and a PE 50 tubing was placed into the i.p. cavity for dexmedetomidine (sedative) infusion. Thereafter, mice were maintained under complete anesthesia with 0.25% isoflurane and 250 μg/kg/h i.p, of dexmedetomidine. Body temperature was maintained at 36°C–37°C using a heating pad and monitored using an MRI-compatible rectal probe throughout the MRI experiments. Heart rate and respiratory rate were continuously monitored throughout the MRI experiments. Dynamic blood oxygenation dependent (BOLD) data were obtained using a Bruker 9.4T/16 magnet (Bruker BioSpin, MA, USA) using a single-shot gradient echo, echo planar imaging (GE-EPI) sequence with the following parameters: TR of 1000 ms, TE of 12 ms, in-plane resolution of 400 × 400 μm and slice thickness of 1000 μm, for a total of 300 images for each run. BOLD time series across all voxels were detrended using a second order fit and bandpass filtered (0.001 to 0.1 Hz) to exclude slow drift of the signal. Images were then registered to a brain template (200 x 200 x 200 um spatial resolution) followed by spatial Gaussian filtering (FWHM=1.5 mm) as previously described (Farina et al., 2021; Sanganahalli et al., 2021).

### Transient microglial ablation

CX3CR1^creET2^-DTA male breeders (C57BL/6J background) that were homozygous for the CX3CR1-Cre-ET2 transgene (Jax #021160) and heterozygous for the floxed diphtheria toxin A gene (R26-eGFP-DTA, (Jax #006331) were mated with Balb/cByj females to generate mixed litters in which half of the pups were CX3CR1-^creET/WT^;DTA^DTA/WT^ (abbreviated as DTA) and the other half are CX3CR1-^creET/WT^;DTA^WT/WT^ (abbreviated as WT). All litters were raised under CTL conditions (i.e. 2 cups of bedding and one nestlet) and at P10 administered tamoxifen i.p. (30 mg/kg) to rapidly ablate microglia. Most pups were weaned at P26 and housed with 2-3 same-sex mice until tested in the contextual fear conditioning at P30-33, although a few pups were sacrificed at different ages to assess microglial elimination or were tested in the open field test at P17. Twenty-four hours after testing contextual freezing, mice were perfused to assess spine density and morphology or glutamatergic spine density. An additional cohort of P30-33 mice were scanned to assess local functional connectivity using rsfMRI.

### Transient chemogenetic activation of microglia

CX3CR1 ^creET2^-Gq-DREADD male breeders (C57BL/6J background) that were homozygous for the CX3CR1-Cre-ET2 transgene (Jax #021160) and heterozygous for the CAG-LSL-Gq-DREADD transgene (Gq-DREADD, Jax# 026220) were mated with Balb/cByj females to generate mixed litters in which half of the pups were CX3CR1-^creET/WT^;Gq-DREADD^Gq/WT^ (abbreviated as Gq) and the other half were CX3CR1-^creET/WT^; Gq-DREADD^WT/WT^ (abbreviated as WT). Litters were randomized to CTL or LB conditions and administered tamoxifen i.p. (30 mg/kg, MP Biomedicals LLC Cat# 156738) at P10 to induce Gq-DREADD expression in microglia. CNO (1 mg/kg, Sigma, Cat#C0832) was then administered i.p. daily on P13, P14, P15, P16, and P17. Some pups were processed at P17, 1hr after the last CNO injection, to assess in vivo phagocytic activity. All other mice were weaned at P26 and housed with 2-3 same-sex mice until tested in the contextual fear conditioning at P30-33. Twenty-four hours after contextual fear conditioning, mice were perfused to assess spine density and morphology or glutamatergic spine density.

### Statistical analysis

The data were carefully screened for inaccuracies, outliers, normality, and homogeneity of variance using SPSS (IBM Corp. version 26) and visualized with GraphPad Prism (iOS, version 10.0). Two-way ANOVA was used to assess the effects of rearing (CTL vs. LB), sex and their interaction on contextual freezing, the maturity index, and glutamatergic synapse density (Fig 1), as well as in vivo phagocytic activity and MEGF10 expression in astrocytes (Fig 6). Significant main effect of rearing or interaction were followed by a preplanned Sidak-post-hoc analysis for each sex. The effects of rearing (CTL vs LB), sex, age (P17 vs P33), and their interaction on microglial-phagocytic activity (Fig 2) were initially examined using a 3-way ANOVA but simplified to a two-way ANOVA focusing on the main effects of rearing and age and their interaction.

**Figure 1.**
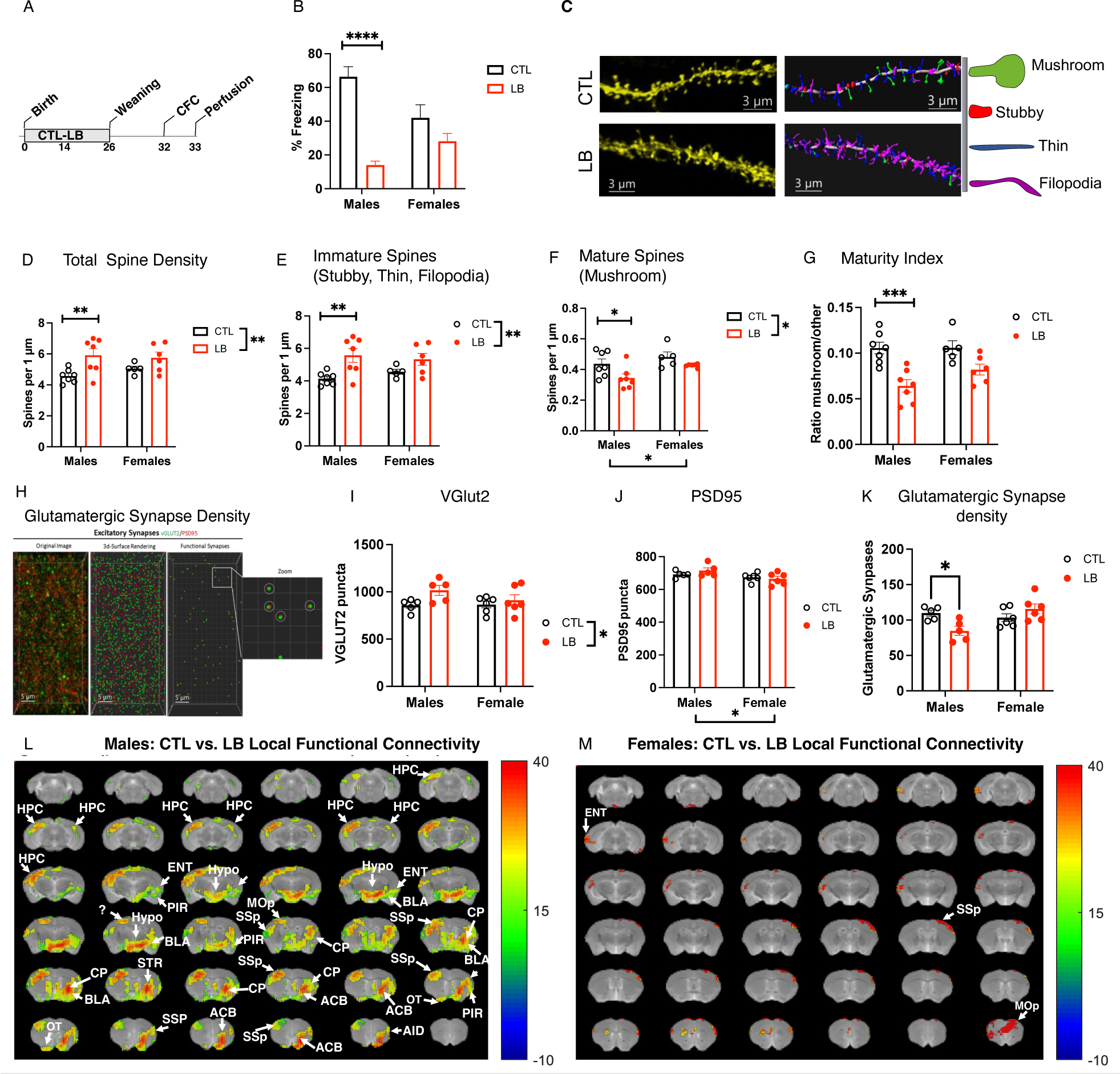
LB Causes Sex-Specific Deficits in Contextual Freezing, Synaptic Maturity, Synaptic Density and Local Functional Connectivity in LB Male Adolescent Mice. (A) Experimental Timeline. (B) Contextual fear conditioning. Rearing: F (1, 42) = 34.44, P< 0.0001, Sex: F (1, 42) = 0.84, P= 0.36, Interaction: F (1, 42) = 11.63 P=0.0014, CTL vs LB- males: P <0.0001, Cohen’s d= 3.5, females: P= 0.16, Cohen’s d= 0.62. N= 11-15 mice per rearing and sex group. (C) DiOlistic images and Imaris models of apical dendrites in the stratum radiatum in adolescent CTL and LB mice. Mushroom spines (green), thin spines (blue), filopodia (magenta). (D) Total Spine Density. Rearing: F (1, 21) = 10.19, P=0.0044, 17p^2^= 0.33, Sex: F (1, 21) = 0.23, P= 0.63, Interaction: F (1, 21) = 0.9, P= 0.33. Sidak post-hoc analysis, CTL vs LB- males: P= 0.0093, Cohen’s d= 1.57. CTL vs LB- females: P= 0.29, Cohen’s d= 0.99. (E) Density of Immature spines. Rearing: F (1, 21) = 11.99, P= 0.0023, 17p^2^= 0.36, Sex F (1, 21) = 0.082, P= 0.77, Interaction: F (1, 21) = 1.135, P= 0.29. Sidak post-hoc analysis, CTL vs LB- males: P= 0.0051, Cohen’s d= 1.71. CTL vs LB- females: P= 0.23, Cohen’s d= 1.07. (F) Mature spines. Rearing: F (1, 21) = 7.827 P=0.011, 17p^2^= 0.27, Sex: F (1, 21) = 5.560 P=0.0282, 17p^2^= 0.21, Interaction: F (1, 21) = 0.4883, P= 0.49. Sidak post-hoc analysis, CTL vs LB- males: P= 0.03, Cohen’s d= 1.22. CTL vs LB- females: P= 0.32, Cohen’s d= 1.16. (G) Maturity Index: Rearing: F (1, 21) = 23.28, P< 0.0005, 17p^2^= 0.53, Sex: F (1, 21) = 1.789, P=0.19, Interaction: F (1, 21) = 1.69, P= 0.21. Sidak post-hoc analysis, CTL vs LB- males: P= 0.0003, Cohen’s d= 2.38. CTL vs LB- females: P= 0.056, Cohen’s d= 1.5. (H) Confocal images and Imaris models used to calculate glutamatergic synapse density in the stratum radiatum. (I) Density of VGlut2 puncta. Rearing: F (1, 18) = 4.816, P= 0.041, 17p^2^= 0.21, Sex: F (1, 18) = 1.049, P= 0.319, Interaction: F (1, 18) = 1.458, P=0.24. Sidak post-hoc analysis, CTL vs LB- males: P= 0.067. CTL vs LB- females: P= 0.95. (J) Density of PSD95 puncta. Rearing: F (1, 18) = 0.3042, P= 0.58, Sex: F (1, 18) = 6.495, P= 0.02, Interaction: F (1, 18) = 1.492, P= 0.24. (K) Density of glutamatergic synapses: Rearing: F (1, 18) = 1.119, P= 0.30, Sex: F (1, 18) = 4.185, P= 0.056, Interaction: F (1, 18) = 9.702, P= 0.006, 17p^2^= 0.35. Sidak post-hoc analysis, CTL vs LB- males: P= 0.022, Cohen’s d= 2.05. CTL vs LB- females: P= 0.29, Cohen’s d= -0.81. (L-M) Effects of rearing on local functional connectivity in males (L) and females (M), FDR < 0.05, local cluster size k> 25 voxels, N= 5-6 mice per rearing and sex group. Abbreviations: ACB- nucleus accumbens, AID- agranular insular area, BLA- basolateral amygdala, CP- caudoputamen, ENT- Entorhinal cortex, HPC- hippocampus, Hypo- hypothalamus, MOp- primary motor area, OT- olfactory tubercle, Pir- piriform area, SSp- primary sensory area.

**Figure 2.**
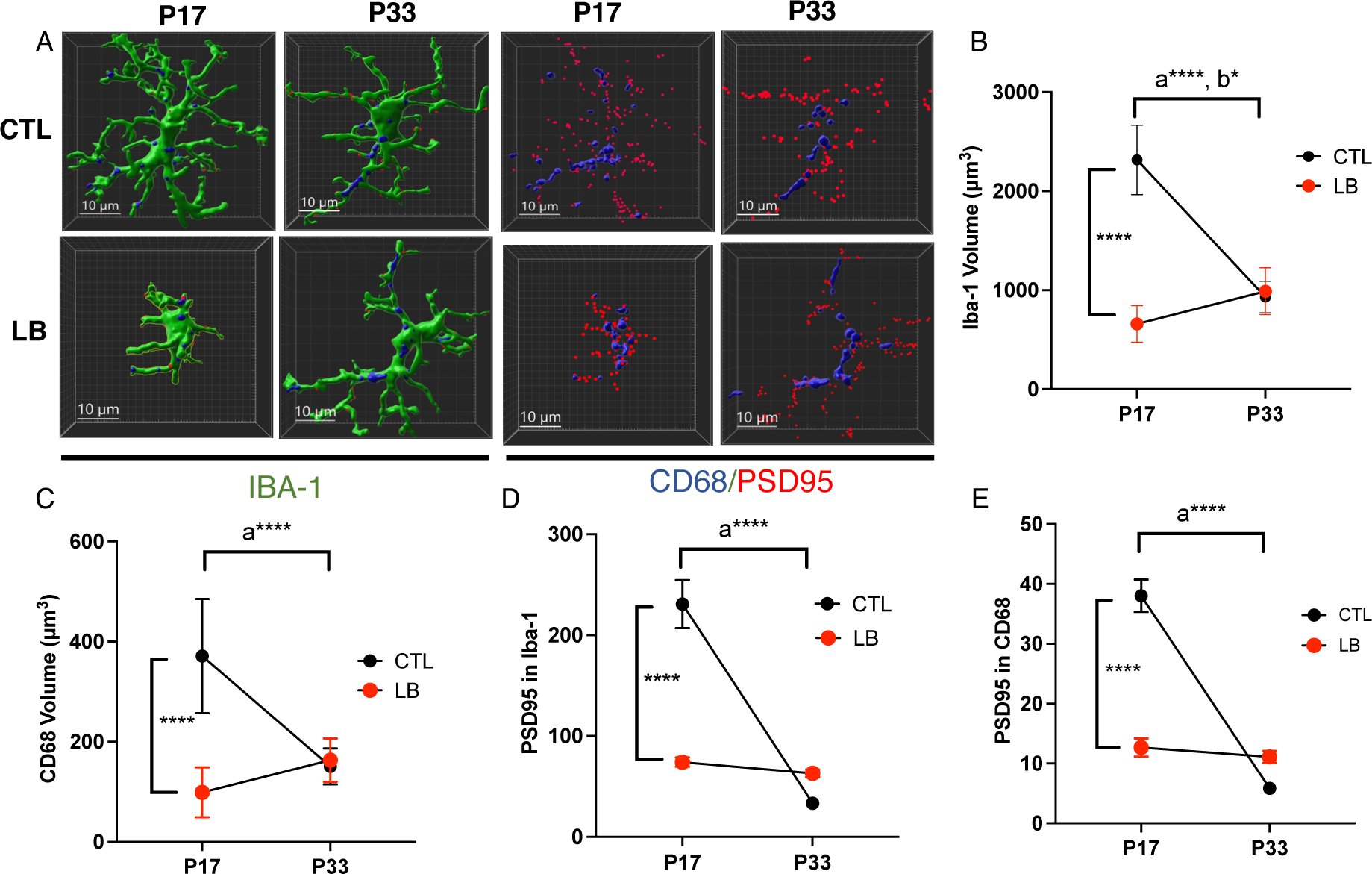
LB Impairs Microglial Mediated Synaptic Pruning at P17 but not P33. (A) Imaris images of microglia (green, left panel) in CTL and LB P17 and P33 mice. CD68 phagosome staining (blue), and PSD95 puncta (red) of the same cells are shown in the right panel. Scale bars 10um. N= 8 mice per rearing and age group, 50% females. (B) Microglial volume. Rearing by age interaction: F (1, 28) = 98.95 P<0.0001, 17p^2^ = 0.78. CTL vs LB P17: P< <0.0001, Cohen’s d= 5.92. CTL vs LB P33: P= 0.63. P17 vs P33 CTL: P< 0.0001, Cohen’s d= 5.09 (shown as lower-case a). P17 vs P33 LB: P= 0.01, Cohen’s d= -1.56 (shown as lower-case b). (C) CD68 volume. Rearing by age interaction: F (1, 28) = 34.82 P<0.0001, 17p^2^ = 0.55. CTL vs LB P17: P< <0.0001 Cohen’s d= 3.1. CTL vs LB P33: P= 0.35. P17 vs P33 CTL: P< <0.0001 Cohen’s d= 2.61 (lower-case a). P17 vs P33 LB: P= 0.25 (D), Number of PSD95 puncta inside microglia. Rearing by age interaction: F (1, 28) = 56.58 P<0.0001, 17p^2^ = 0.67. CTL vs LB P17: P< <0.000, Cohen’s d= 3.23. CTL vs LB P3: P= 0.98. P17 vs P33 CTL P< <0.0001, Cohen’s d= 4.1 (shown as a), P17 vs P33 LB: P= 0.92. (E) Number of PSD95 puncta inside CD68 phagosome. Rearing by age interaction: F (1, 28) = 86.90, P<0.0001, 17p^2^ = 0.75. CTL vs LB P17: P< <0.0001, Cohen’s d= 4.1. CTL vs LB P33: P= 0.18. P17 vs P33 CTL: P< <0.0001, Cohen’s d= 5.88 (lower-case a). P17 vs P33 LB: P= 0.98.

Significant interaction between rearing and age was followed by a preplanned Tukey-HSD post-hoc comparison across all groups. For microglial ablation, a two-way ANOVA was used to assess the effects of genotype (WT vs DTA), sex, and their interaction on contextual freezing, the maturity index, and glutamatergic synapse density with significant main effects of genotype or interaction followed by preplanned Sidak-post-hoc analyses for each sex. Three-way ANOVA was initially used to assess the effects rearing (CTL vs LB), sex, and genotype (WT vs. Gq) on microglial phagocytic activity in-vivo in P17 pups. However, since similar outcomes were seen in males and females, this was simplified to a two-way ANOVA focusing on the effects of rearing and genotype with significant interaction followed by Tukey-HSD post-hoc comparisons across all groups (Fig 4). Three-way ANOVA was also used to assess the effects of rearing (CTL vs LB), sex, and genotype (WT vs. Gq) on contextual freezing in P33 adolescent mice. However, since there was a significant interaction between rearing and sex, we conducted separate two-way analyses in males and females. Significant effects of rearing or interaction were followed by preplanned Tukey-HSD post-hoc comparisons across all groups. Local functional connectivity maps were calculated for each voxel using threshold correlation (Tc) > 0.6, a signal-to-noise ratio (TSNR> 0.5), and distance < 1 mm in individual space and normalized to a z score with the Fisher r to z transformation. Significance was assessed using same-sex Student’s t-test comparisons for different rearing conditions with Benjamini-Hochberg correction for multiple comparisons with a false discovery rate (FDR) < 0.05 and local cluster size of k> 25 voxels.

**Figure 3.**
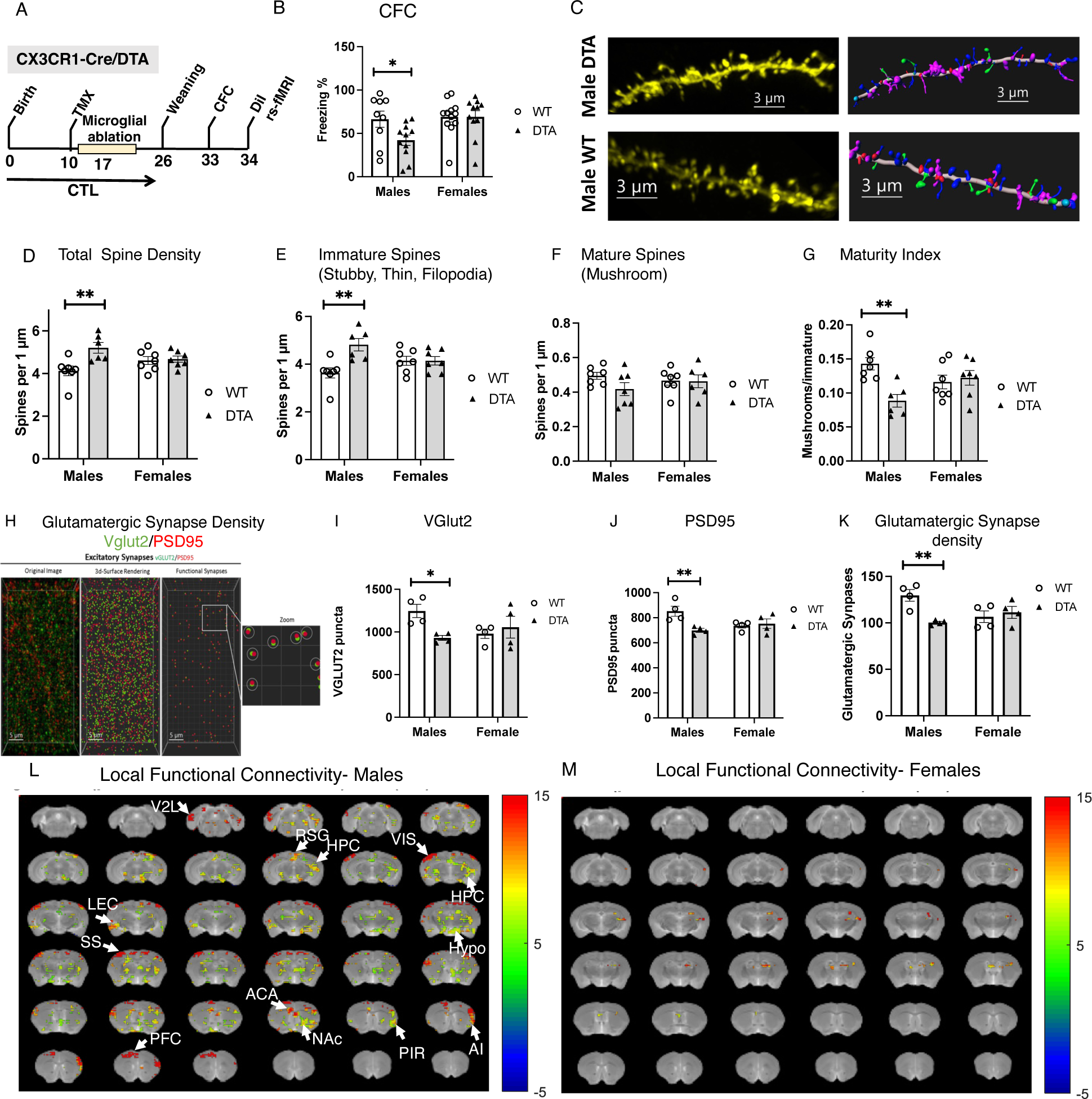
Transient Ablation of Microglia Induces Similar Sex-Specific Changes Observed in LB Adolescent Mice. (A) Experimental Timeline. B. Contextual fear conditioning. Genotype: F (1, 39) = 3.00, P=0.09, Sex: F (1, 39) = 4.49, P=0.04, Interaction: F (1, 39) = 2.98, P= 0.092. Post-hoc CTL vs LB- males: P= 0.046. CTL vs LB- females: P >0.99. (B) Confocal images and Imaris models of DiOlistic labeling of apical dendrites in the stratum radiatum in adolescent WT and DTA male mice. Mushroom spines (green), thin spines (blue), filopodia (magenta). (C) Total spine density. Genotype: F (1, 23) = 8.097, P= 0.0092, Sex: F (1, 23) = 0.012, P= 0.91, Interaction: F (1, 23) = 6.374, P=0.019. Post-hoc CTL vs LB- males: P= 0.0022. CTL vs LB- females: P= 0.97. (D) Immature spines. Genotype: F (1, 23) = 8.020, P=0.0094, Sex: F (1, 23) = 0.1401, P=0.712, Interaction: F (1, 23) = 8.05, P= 0.0093. Post-hoc CTL vs LB- males: P= 0.0013. CTL vs LB- females: P >0.99. (F). Mature Spines. Genotype: F (1, 23) = 1.788, P= 0.19, Sex: F (1, 23) = 0.078, P= 0.78, Interaction: F (1, 23) = 1.41, P= 0.25. (G) Maturity Index. Genotype: F (1, 23) = 5.99, P= 0.022, Sex: F (1, 23) = 0.12, P= 0.731, Interaction: F (1, 23) = 9.425, P=0.0054. Post-hoc CTL vs LB- males: P= 0.0017. CTL vs LB- females: P= 0.88. (H) Confocal images and Imaris models of VGlut2 and PSD95 puncta in the stratum radiatum. (I) VGlut2 density. Genotype: F (1, 12) = 2.19, P= 0.16, Sex: F (1, 12) = 0.76, P= 0.39, Interaction: F (1, 12) = 5.874, P=0.032. Post-hoc CTL vs LB- males: P= 0.035. CTL vs LB- females: P> 0.99. (J) PSD95 density. Genotype: F (1, 12) = 5.09, P= 0.043, Sex: F (1, 12) = 1.09, P= 0.32, Interaction: F (1, 12) = 7.827, P=0.016. Post-hoc CTL vs LB- males: P= 0.0077, Females: P> 0.99. (K) Glutamatergic Synapse Density. Genotype: F (1, 12) = 5.014, P=0.04, Sex: F (1, 12) = 1.19, P= 0.29, Interaction: F (1, 12) = 9.58, P= 0.0093. Post-hoc CTL vs LB- males: P= 0.0053, females: P> 0.99. (L-M) Local functional connectivity. Local functional connectivity maps of WT vs DTA males (L) and females (M), FDR < 0.05, local cluster size k> 25 voxels, N= 6 mice per rearing and sex group (red-yellow colors indicate reduced connectivity in DTA compared to WT mice). Abbreviations: ACA- Anterior Cingulate Area, AI- Anterior Insular area, V2L secondary visual lateral Cortex, RSG- retrosplenial granular area, HPC- hippocampus, VIS- visual cortex, LEC- lateral entorhinal cortex, Hypo- hypothalamus, NAc- nucleus accumbens, PIR- piriform area, SS- Somatosensory area, PFC- medial prefrontal cortex.

**Figure 4.**
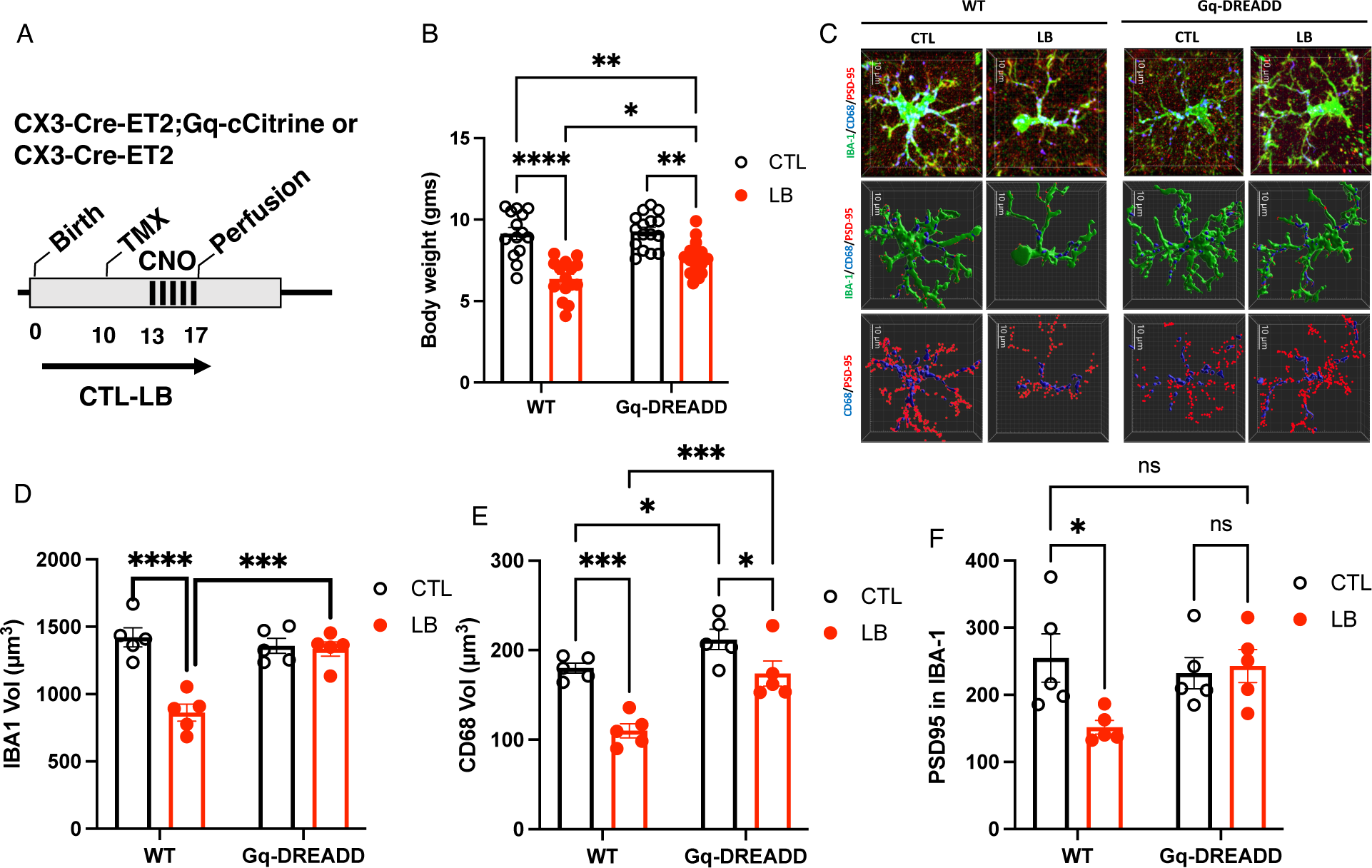
Chemogenetic Activation of Microglia Restores Normal Phagocytic Activity in P17 LB Mice. (A) Experimental Timeline. (B) Effects of rearing and genotype on body weight. Rearing: F (1, 57) = 55.87, P< 0.0001, Genotype: F (1, 57) = 5.714, P= 0.020. Interaction: F (1, 57) = 3.86, P= 0.054. Tukey-HSD post-hoc CTL-WT vs LB-WT: P <0.0001. CTL-Gq vs LB-Gq: P= 0.0010, LB-WT vs LB-Gq: P= 0.015. CTL-WT vs LB-Gq: P= 0.0041. Half of the animals are females. (C) Confocal and Imaris images of IBA1 (green), CD68 (blue), and PSD95 puncta (red) staining of microglia located in the stratum radiatum of P17 mice administered daily CNO injections from P13-17. (D) Effects of rearing and genotype on microglial volume. Rearing: F (1, 16) = 22.88, P= 0.0002, Genotype: F (1, 16) = 11.25, P= 0.0040. Interaction: F (1, 16) = 19.35, P= 0.0004. Tukey-HSD post-hoc CTL-WT vs LB-WT: P <0.0001. CTL-Gq vs LB-Gq: P >0.99, LB-WT vs LB-Gq: P= 0.0003. CTL-WT vs LB-Gq: P >0.99. Half of the animals are females. (E) Effects of rearing and genotype on CD68 volume. Rearing: F (1, 16) = 27.92, P< 0.0001, Genotype: F (1, 16) = 21.93, P= 0.0002. Interaction: F (1, 16) = 2.42, P= 0.14. Tukey-HSD post-hoc CTL-WT vs LB-WT: P= 0.0002. CTL-Gq vs LB-Gq: P= 0.0180, LB-WT vs LB-Gq: P= 0.0004. CTL-WT vs LB-Gq: P= 0.68. Half of the animals are females. (F) Effects of rearing and genotype on PSD95 engulfed by microglia. Rearing: F (1, 16) = 3.35, P= 0.086, Genotype: F (1, 16) = 1.84, P=0.19. Interaction: F (1, 16) = 5.033, P= 0.039. Tukey-HSD post-hoc CTL-WT vs LB-WT: P= 0.048. CTL-Gq vs LB-Gq: P= 0.99, LB-WT vs LB-Gq: P= 0.09. CTL-WT vs LB-Gq: P= 0.99. Half of the animals are females.

## RESULTS

### LB Causes Sex-Specific Structural and Functional Synaptic Deficits in Adolescent Male Mice

We have recently demonstrated that adolescent male LB mice exhibit more severe deficits in contextual fear conditioning compared to female LB mice (Islam et al., 2023). Here, we replicated these findings using an independent cohort of mice (Figure 1A-B). At the completion of the behavioral testing, mice were processed to evaluate the effects of rearing and sex on spine density and morphology using DiOlistic labeling (Figure 1C and Figure S1A). A 2 x 2 ANOVA revealed a significant effect of rearing on total spine density, with no significant main effect of sex or interaction (Figure 1D). Sidak post-hoc analysis revealed a significant increase in total spine density in males but not females (Figure 1D). The increase in spine density was due to a higher number of immature spines, which was again significant in males but not females (Figure 1E). A 2 x 2 ANOVA of the density of mature (mushroom) spines found significant effects of rearing and sex, but no significant interaction. Sidak post-hoc analysis indicated a significant decrease in the density of mushroom spines in males but not in females (Figure 1F).

We defined the “maturity index” as the ratio between mature mushroom spines and immature spines (e.g., filopodia, thin, stubby; Figure S1B) and found a significant effect of rearing, but no significant effect of sex or interaction (Figure 1G). Post-hoc analysis in males found a highly significant reduction in maturity index while only a trend was observed in LB females (Figure 1G). There was also a highly significant correlation between the maturity index and freezing behavior (r= 0.43, P= 0.007, Figure S1C).

Next, we evaluated the impact of rearing and sex on the density of glutamatergic synapses in the stratum radiatum (Figure 1H). For the presynaptic marker VGlut2, we found a significant effect of rearing, but no significant effects of sex or interaction. A preplanned Sidak post-hoc analysis revealed a non-significant trend for increased VGlut2 density in males and no difference in females (Figure 1I). For the excitatory postsynaptic marker PSD95, there was a significant effect of sex, with a small increase observed in males, but no significant effects of rearing or interaction (Figure 1J). Consistent with the DiOlistic results, we found a significant interaction between LB and sex for the number of functional glutamatergic synapses. This was due to a decrease in excitatory synapses in LB males but not in LB females (Figure 1K).

To investigate the impact of LB and sex on local functional connectivity throughout the entire brain, we adopted an MRI-based method previously employed in human subjects (Tomasi and Volkow, 2010) and applied it to rodents. This approach generates whole-brain voxel maps corrected for multiple comparisons and unbiasedly identify brain regions with abnormal local functional connectivity. Adolescent male LB mice showed reduced local connectivity in multiple brain regions, including the hippocampus, entorhinal cortex, striatum, basolateral amygdala, hypothalamus, and the nucleus accumbens (Figure 1L). In contrast, there was no reduction in local functional connectivity in the hippocampus of LB females with only few brain regions showing significantly lower local functional connectivity (Figure 1M). These findings reveal novel sex differences in the impact of LB on contextual fear conditioning, maturity index, glutamatergic synapse density, and local functional connectivity in the hippocampus of adolescent mice.

### LB Transiently Impairs Microglial Function in the Developing Hippocampus

LB impairs microglial ramification and their ability to engulf synaptic material at P17, an age when the hippocampus undergoes intense synaptic pruning (Dayananda et al., 2022). To determine if these changes persisted in P33 adolescent mice and whether they differentially impacted males, we evaluated the effects of rearing, sex, and age in P17 and P33 mice. No significant main effect or interaction with sex were found, so the data for males and females were combined. A 2 x 2 ANOVA found a significant interaction between rearing and age for microglial volume. Post-hoc analysis comparing CTL and LB P17 pups replicated our previous work showing a significant reduction in microglial volume (Fig 2 A & B), CD68 volume (Fig 2 A & C), the number of PSD96 engulfed by microglia (Fig 2 A & D), and the number of PSD95 inside CD68 (Fig 2 A & D), all with large effect sizes. No significant differences between CTL and LB survived multiple comparisons at P33 when using the entire set of data. A 2 x 2 ANOVA examining the effects of rearing and sex at P33, revealed a significant increase in the number of PSD95 puncta inside microglia (Fig S2C) and inside CD68 (Fig S2D) in LB male and female mice, suggesting a possible compensatory microglial response at this age.

CTL mice exhibited a dramatic reduction in microglial volume, phagosome size, and the number of PSD95 engulfed by microglia at P33 compared to P17 (Figure 2). This is consistent with previous work showing that synaptic pruning reaches its peak in the hippocampus during the second and third weeks of life (Filipello et al., 2018; Scott-Hewitt et al., 2020; Zhan et al., 2014). In contrast, LB microglia showed a small but significant increase in volume at P33 compared to P17 (Figure 2B). However, there were no significant differences in CD68 size, PSD95 puncta inside microglia, or PSD95 puncta inside CD68 between P17 and P33 (Figure 2 C-E). Together, these findings indicate that LB impairs the ability of microglia to engulf synaptic material at P17. These deficits are no longer observed at P33 when microglial phagocytic activity is significantly lower.

### Transient Ablation of Microglia Induces Sex-Specific Changes in the Hippocampus that Resemble Abnormalities Observed in LB Mice

Our data indicate that LB transiently impairs microglial phagocytic activity during the second and third weeks of life when the hippocampus undergoes intense synaptic pruning. These changes are associated with sex-specific impairments in contextual fear conditioning, maturity index, glutamatergic synaptic densities, and local functional connectivity. To test whether transient perturbation in microglial activity can induce similar sex-specific changes, we developed a method to transiently ablate microglia during the 2^nd^-3^rd^ weeks of life and tested the impact of this manipulation on contextual fear conditioning, maturity index, glutamatergic synapse density, and local functional connectivity in adolescent male and female mice. To transiently ablate microglia, we mated CX3CR1-^Cre-ET2/Cre-ET2^; ROSA26-eGFP-^DTA^/^wt^ male mice with Balb/cByj females to generate mixed litters that are either CX3CR1-^Cre-ET2/WT^; ROSA26-eGFP-^DTA^/^wt^ (DTA) or CX3CR1-^Cre-ET2/WT^; ROSA26-eGFP-^wt^/^wt^ (WT) littermates (Fig 3A-B and Fig S2A).

A single injection of tamoxifen at P10 induced rapid elimination of microglia in DTA, but not WT littermates with no signs of toxicity (Figure S3). By P21, the number and the morphology of microglia in the hippocampus of DTA mice were indistinguishable from WT littermates (Fig S3), consistent with previous work showing that a small pool of microglia survives the initial ablation and proliferates to restore normal number of microglia within 1-2 weeks (Bruttger et al., 2015; Nelson and Lenz, 2017; Rice et al., 2015; Schalbetter et al., 2022). Next, we tested the impact of transiently ablating microglia on contextual fear conditioning in adolescent mice raised under CTL condition (Fig 3A-B). Similar to findings with LB mice, transient ablation of microglia reduced contextual freezing in male DTA, but not female DTA mice (Figure 3B). DTA male mice, but not females showed an increase in the total number of spines (Figure 3 C-D) due to the retention of immature spines (Figure 3E). The mean number of mature mushroom spines was reduced in male DTA mice, but this did not reach significance (Figure 3F). The maturity index was reduced in DTA males, but not female adolescent mice (Figure 3G) and as with LB adolescent mice (Figure S1B), there was a significant correlation between the maturity index and freezing behavior (r= 0.74, P= 0.0027, Figure S1).

Unlike outcomes seen in adolescent LB mice, transient ablation of microglia decreased the density of VGlut2 (Figure 3 H & I) and PSD95 (Figure 3 H & J) in males, but not females. However, the density of functional glutamatergic synapses was reduced in DTA males but not in female DTA (Figure 3 K), further highlighting similarities with LB mice (see Figure 1K). DTA male mice had reduced local functional connectivity compared to WT littermates across multiple brain regions, including the hippocampus (Figure 3L), changes that were not seen in DTA females (Figure 3M). Together, these findings indicate that transient ablation of microglial during the second and third weeks of life can replicate several of the sex-specific changes seen in the hippocampus of LB adolescent mice.

### Chemogenetic Activation of Microglia Restores Phagocytic Activity in P17 LB Mice

To test whether chemogenetic activation of microglia during the second and third weeks of life can normalize microglial-mediated synaptic pruning in 17-day-old LB pups, we mated CX3CR1- ^Cre-ET2^/ ^Cre-ET2^/; CAG-LSL-Gq-DREADD^Gq/WT^ males to Balb/cByj females to generate mixed litters that are either CX3CR1-^Cre-ET2^/ ^WT^; CAG-LSL-Gq-DREADD^Gq/WT^ (abbreviated as Gq) or CX3CR1- ^Cre-ET2^/ ^WT^; CAG-LSL-Gq-DREADD^WT/WT^ (abbreviated as WT) and randomized them to either CTL or LB conditions. At P10 tamoxifen was administered to express the Gq-DREADD followed by five daily injections of CNO from postnatal day 13 to 17. Pups were perfused 60 minutes after the last CNO injection to assess microglial morphology and phagocytic activity (Figure 4 A-B).

All microglia from Gq mice expressed the HA tag, and no HA tag was expressed in microglia from WT mice (Figure S4). No sex differences or interaction with sex were found for any tested variables and therefore data from male and female mice were combined to determine main effects of rearing, genotype, and their interaction. For body weight, there was a significant main effect of rearing and genotype, with a trend for a significant interaction. Tukey-HSD post hoc analysis confirmed our previous work showing a reduction in body weight in LB pups. This was highly significant for CTL-WT vs LB-WT mice and to a lesser degree, although still significant, for CTL-Gq vs LB-Gq mice (Figure 4B). Interestingly, chemogenetic activation of microglia increased the body weight of LB (LB-WT vs LB-Gq) but not CTL mice (CTL-WT vs CTL-Gq) indicating that abnormal microglial activity contributes to the reduction in LB body weight. Chemogenetic activation increased the volume of microglia, CD68 size, and the number of PSD95 puncta inside microglia in LB-Gq pups to levels observed in CTL-WT pups with similar outcomes seen in males and females (Figure 4C-F). These findings indicate that chemogenetic activation of microglia during the second and third weeks of life normalizes microglia-mediated synaptic pruning in the developing hippocampus of mice exposed to LB.

### Chemogenetic Activation of Microglia During the 2^nd^ and 3^rd^ Weeks of Life Restores Normal Hippocampal Function in Adolescent LB males

Next, we tested whether chemogenetic activation of microglia from P13-17 can reverse the contextual fear conditioning deficits observed in adolescent LB mice (Figure 5A). A three-way ANOVA revealed a significant interaction between rearing, sex, and genotype (F (1, 107) = 5.643, P= 0.019), prompting us to conduct separate analyses in males and females. Males exhibited a significant interaction between rearing and genotype due to reduced freezing in LB- WT compared to CTL-WT and LB-Gq, with no significant differences between LB-Gq and LB- WT or CTL-WT mice (Figure 5B). As expected, no significant effects of rearing, genotype, and interaction were seen in females (Figure 5C). We then focused on the effects of chemogenetic activation on spine density and morphology in CA1 apical dendrites (Figure 5D). LB increased the total spine density in WT mice, but not in Gq mice, with no significant difference between CTL-WT and LB-Gq mice (Figure 5E), with similar outcomes observed for the number of immature spines (Figure 5F). Chemogenetic activation also normalized the number of mature spines in LB male mice (Figure 5G) and significantly increased the maturity index in LB-Gq mice to levels observed in CTL-Gq mice, although it did not fully restore the maturity index to CTL- WT levels (Figure 5F). The maturity index was again highly correlated with freezing behavior (r= 0.87, P= 0.001). LB increased the number of the presynaptic Vglut2 puncta in WT but not Gq mice (Figure 5 I & J), while levels of PSD95 were unexpectedly higher in LB-Gq compared to CTL-WT or CTL-Gq mice (Figure 5K). Chemogenetic activation completely restored the density of functional glutamatergic synapses in the stratum radiatum (Figure 5L).

**Fig 5.**
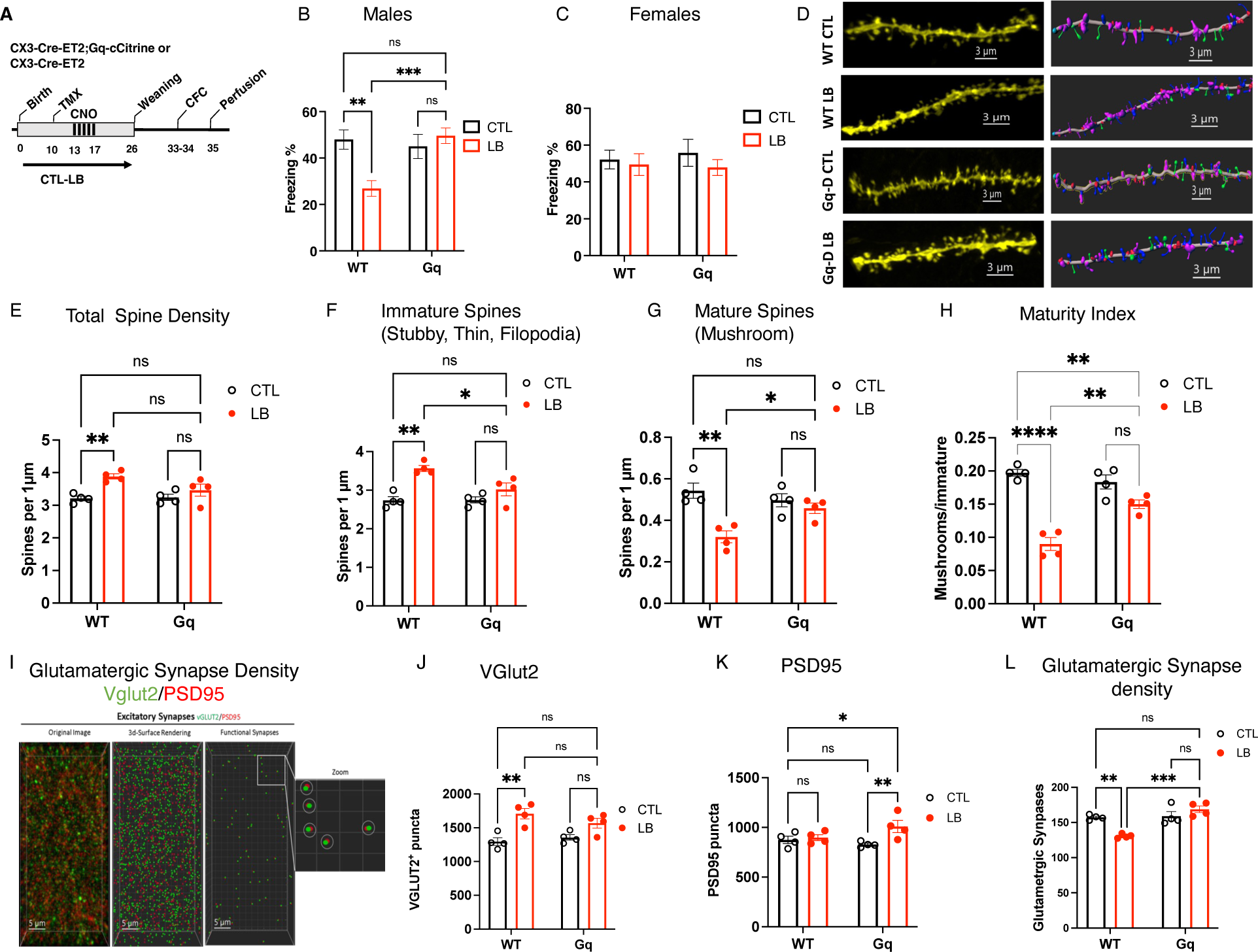
Chemogenetic Activation of Microglia During the Second and Third Weeks of Life Normalizes Contextual Fear Conditioning and Synaptic Abnormalities in Adolescent LB Males. (A) Experimental timeline. (B-C) contextual fear conditioning. (B) Males, n=12-19 mice per group. Rearing: F (1, 58) = 4.28, P= 0.043, Genotype F (1, 58) = 6.197, P=0.016, Interaction: F (1, 58) = 10.44, P=0.0020. CTL vs LB WT: P= 0.0011, CTL vs LB Gq: 0.97. LB-WT vs LB-Gq P= 0.0003, CTL-WT vs LB-Gq P= 0.99. (C) Females, n=11-18 mice per group. Rearing: F (1, 48) = 0.91, P= 0.35, Genotype: F (1, 48) = 0.033, P= 0.85, Interaction: F (1, 48) = 0.22, P= 0.64. (D) Confocal images and Imaris models of DiOlistic labeling of apical dendrites in the stratum radiatum of adolescent male mice. Mushroom spines (green), thin spines (blue), filopodia (magenta). (E) Total spine density. Rearing: F (1, 12) = 2.66, P= 0.12, Genotype: F (1, 12) = 14.41,P= 0.0025, Interaction: F (1, 12) = 3.66, P= 0.07. CTL-WT vs LB-WT: P= 0.0077, CTL-Gq vs LB-Gq: P= 0.56. LB-WT vs LB-Gq P= 0.11, CTL-WT vs LB-Gq P= 0.45. (F) Immature spines. Rearing: F (1, 12) = 5.74, P= 0.0338, Genotype: F (1, 12) = 24.36, P= 0.0003, Interaction: F (1, 12) = 6.172, P=0.029. CTL-WT vs LB-WT: P= 0.0012, CTL-Gq vs LB- Gq: P= 0.49. LB-WT vs LB-Gq P= 0.028, CTL-WT vs LB-Gq P= 0.46. (G) Mature spines. Rearing: F (1, 12) = 2.165, P= 0.16, Genotype: F (1, 12) = 17.81, P= 0.0012, Interaction: F (1, 12) = 8.84, P= 0.0116. CTL-WT vs LB-WT: P= 0.0013, CTL-Gq vs LB-Gq: P= 0.81. LB-WT vs LB-Gq P= 0.037, CTL-WT vs LB-Gq P= 0.26. (H) Maturity Index. Rearing: F (1, 12) = 7.79, P= 0.016, Genotype: F (1, 12) = 72.83, P< 0.0001, Interaction: F (1, 12) = 20.09, P= 0.0008. CTL-WT vs LB-WT: P <0.0001, CTL-Gq vs LB-Gq: P= 0.082. LB-WT vs LB-Gq P= 0.0015, CTL-WT vs LB-Gq P= 0.0094. (I) Confocal images and Imaris models of VGlut2 and PSD95 puncta in the stratum radiatum. (J) VGlut2 density. Rearing: F (1, 12) = 24.22, P= 0.0004, Genotype: F (1, 12) = 0.39, P= 0.5437, Interaction: F (1, 12) = 2.47, P= 0.14. CTL-WT vs LB-WT: P= 0.0037, CTL-Gq vs LB-Gq: P= 0.21. LB-WT vs LB-Gq: P= 0.87, CTL-WT vs LB-Gq: P= 0.062. (K) PSD95 density. Rearing: F (1, 12) = 6.676 P=0.024, Genotype: F (1, 12) = 0.68, P= 0.42, Interaction: F (1, 12) = 4.068, P=0.067. CTL-WT vs LB WT: P= 0.69, CTL-Gq vs LB-Gq: P= 0.0069. LB-WT vs LB-Gq: P= 0.067, CTL-WT vs LB-Gq: P= 0.032. (L) Glutamatergic synapse density. Rearing: F (1, 12) = 3.924 P=0.071, Genotype: F (1, 12) = 22.36, P= 0.0005, Interaction: F (1, 12) = 18.61 P=0.0010. CTL-WT vs LB-WT: P= 0.0047, CTL- Gq vs LB-Gq: P= 0.75. LB-WT vs LB-Gq: P= 0.0002, CTL-WT vs LB-Gq: P= 0.45.

### LB Increases MEGF10 and Phagocytic Activity in Females’ Astrocytes

Our data suggest that during the second and third weeks of life, female LB mice are somehow able to compensate for deficits in microglial-mediated synaptic pruning. Given that astrocytes also play an important role in synaptic pruning and, in some cases, can compensate for abnormal microglia-mediated synaptic pruning (Chung et al., 2013; Damisah et al., 2020; Konishi et al., 2020; Lee et al., 2021; Perez-Catalan et al., 2021), we tested whether 17-day-old female LB mice can upregulate phagocytic activity in astrocytes (Figure S5A-B). No significant effects of rearing, sex, or their interaction were found for the number of GFAP-positive astrocytes in the stratum radiatum (Figure S5C). However, there was a significant interaction between rearing and sex for the number of PSD95 puncta per astrocyte and the density of PSD95 inside astrocytes. Post-hoc analysis revealed that this interaction was due to increase in PSD95 puncta in LB females but not in LB males (Figure S5 D-E). Previous studies have demonstrated that phagocytic activity in astrocytes is regulated by the MerTK and MEGF10 receptors (Chung et al., 2013; Lee et al., 2021). We therefore teste the levels of these receptors in LB male and female mice. No MerTK, staining was observed in GFAP-positive astrocytes in the stratum radiatum at this age. However, MEGF10 staining was clearly localized within GFAP-positive cells (Figure 6). We therefore assessed the effects of rearing and sex on MEGF10 and PSD95 puncta in astrocytes using a second cohort of 17-day-old pups (Figure 6A). There was a significant effect of rearing, but no significant effects of sex or interaction for the volume of GFAP-positive astrocytes located in the stratum radiatum (Figure 6B). We replicated our initial findings, showing a significant increase in the number of PSD95 inside astrocytes in LB females, but not LB males (Figure 6C and Fig S5). Additionally, this sex-specific effect was accompanied by an increase in MEGF10 staining in astrocytes from LB females, but not LB males (Figure 6D). Next, we investigated whether similar sex-specific changes were seen in 17-day old mice exposed to transient ablation of microglia. Similar to outcomes seen in LB mice, transient ablation of microglia led to an increase in the size of GFAP-positive astrocytes in female subjects, but not in males (Fig S6A-B). However, unlike the outcomes in LB, microglial ablation increased the number of PSD95 puncta and MEGF10 staining in both males and females (Figure S6 C & D).

**Fig 6.**
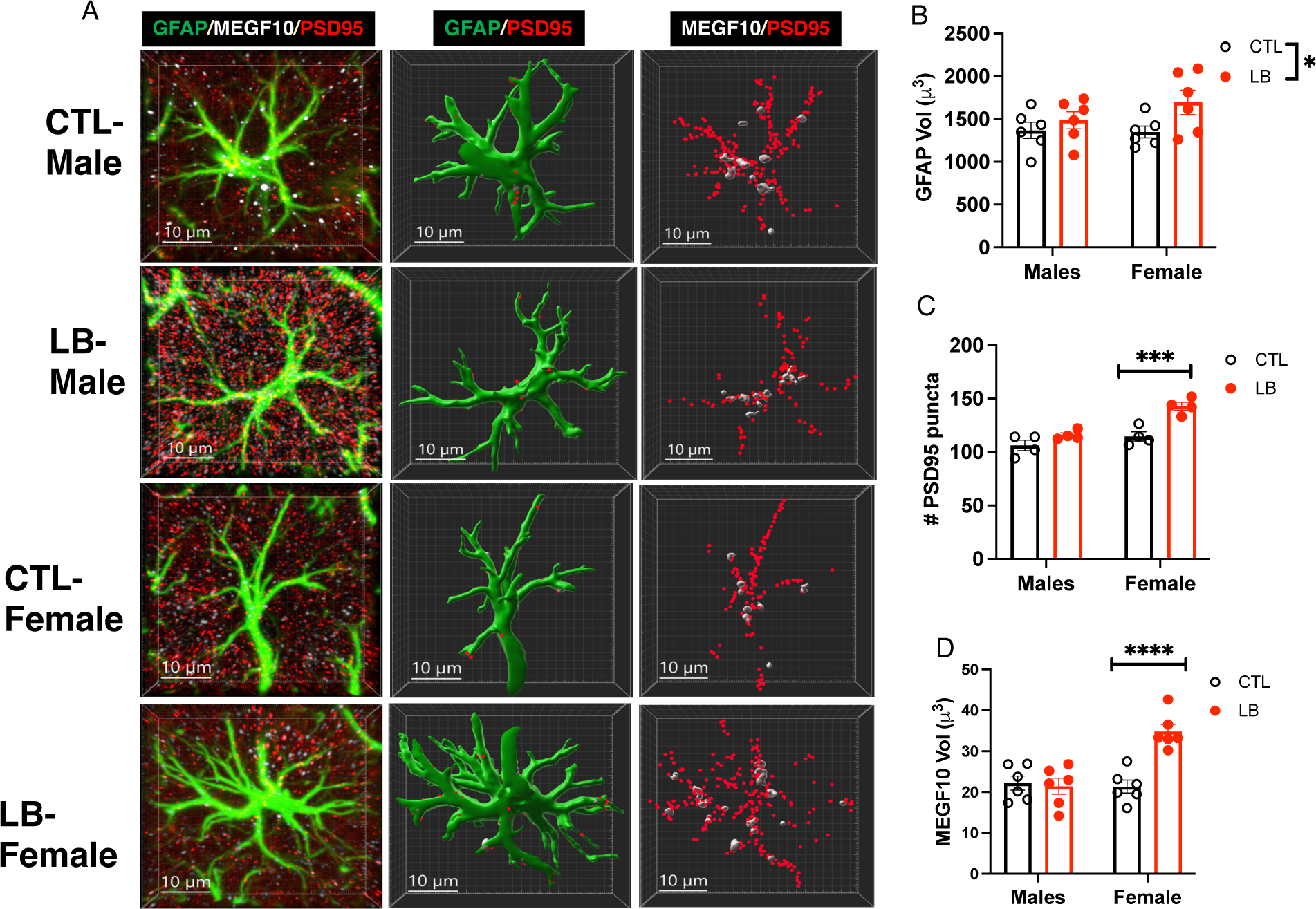
LB Increases MEGF10 and Phagocytic Activity in Female Astrocytes. (A) Confocal images and Imaris models of GFAP-positive astrocytes in the stratum radiatum of 17-day old CTL and LB pups. GFAP (green), PSD95 puncta (red), MEGF10 (white). Scale bar 10μm. (B) Astrocyte cell volume. Rearing: F (1, 20) = 4.916, P= 0.038, Sex: F (1, 20) = 0.82, P= 0.38, Interaction: F (1, 20) = 1.196, P= 0.29. CTL vs LB Males: P= 0.68, CTL vs LB Females: P= 0.059. (C) PSD95 puncta inside astrocyte. Rearing: F (1, 12) = 23.02, P= 0.0004, Sex: F (1, 12) = 20.61, P= 0.0007, Interaction: F (1, 12) = 5.583, P= 0.036, CTL vs LB Males: P= 0.2092, CTL vs LB Females: P= 0.0006. (D) MEGF10 staining inside astrocyte. Rearing: F (1, 20) = 12.92, P= 0.0018, Sex: F (1, 20) = 12.52, P= 0.0021, Interaction: F (1, 20) = 16.19, P=0.0007, CTL vs LB Males: P= 0.76, CTL vs LB Females: P< 0.0001.

## DISCUSSION

One of the most consistent findings in individuals exposed to ELA is abnormal hippocampal function, which seems to be more pronounced in men than in women (Garvin and Bolton, 2022; Teicher and Samson, 2016; White and Kaffman, 2019). However, little is currently known about the nature of these sex-specific structural and functional changes in the hippocampus and their impact on cognition. Furthermore, it is unclear which forms of ELA cause sexually dimorphic changes and what underlying mechanisms are responsible for the different outcomes in males and females. These are difficult questions to address in humans, but they can be investigated in rodent models of ELA and further validated with imaging techniques that can be used across species (Kaffman et al., 2019). However, most of the research on rodents to date has focused on changes in adult males. Therefore, additional studies are needed to examine changes in both sexes, especially during adolescence when sex differences might be more prominent (Gershon et al., 2008; White and Kaffman, 2019). Moreover, most studies have examined the effects of ELA on neurons, with relatively little attention paid to the possible contribution of glial cells. In particular, the role of microglia in inducing sexually dimorphic changes in ELA needs further investigation, considering the crucial role of microglia in establishing sexually dimorphic changes in the developing brain (VanRyzin et al., 2020) and sex-specific responses to pain, obesity (Dorfman et al., 2017), and ischemia (Villa et al., 2018). This study presents several novel findings about the role of glial cells in programming sexually dimorphic changes in the hippocampus of adolescent mice exposed to LB, a commonly used mouse model of ELA.

### LB causes sex-specific deficits in the stratum radiatum and contextual fear conditioning in adolescent mice

Our work supports the emerging notion that LB causes more severe hippocampal-dependent deficits in males compared to females (Bath et al., 2017; Islam et al., 2023; Naninck et al., 2015). Naninck and colleagues provided evidence indicating that these sex differences were due to more severe deficits in adult neurogenesis in males (Naninck et al., 2015), but these findings have not been replicated by others (Bath et al., 2017; Youssef et al., 2019). We have recently shown that LB causes significant deficits in contextual fear conditioning in adolescent male, but not female, mice (Islam et al., 2023). LB reduced hippocampal volume in both adolescent male and female mice, indicating that LB female mice are also affected by LB and that volumetric changes are unlikely to explain the sex-specific abnormalities in contextual fear conditioning (Islam et al., 2023). Instead, we found that the connectivity between the entorhinal cortex and the dorsal hippocampus was reduced fourfold in LB males but not in female mice. Given the critical role that these projections play in mediating contextual fear conditioning we proposed that changes in connectivity contribute to sex differences in hippocampal function (Islam et al., 2023).

Here we examine the effects of rearing and sex on spine density and morphology of CA1 pyramidal neurons apical dendrites located in the stratum radiatum. These apical dendrites receive input from CA3 pyramidal neurons via the Schaffer collaterals and play a critical role in hippocampal mediated tasks such as contextual fear conditioning (Astudillo et al., 2020; Basu et al., 2016; Li et al., 2005; Sacchetti et al., 2001; Sun et al., 2016). Using DiOlistic labeling we found that LB male, but not LB female, mice show higher levels of immature spines (e.g., stubby, thin, filopodia) and low levels of mature mushroom spines. We found that the maturity index is significantly reduced in male LB mice, but not females, and is highly correlated with contextual freezing. Sex-specific deficits were also observed in the density of glutamatergic synapse density in the stratum radiatum and local functional connectivity using rsfMRI. Together, these findings uncover novel differences in synaptic function in the hippocampus of adolescent male and female mice exposed to LB providing a likely explanation for the sex differences seen in contextual fear conditioning. We provide evidence that alterations in glial mediated synaptic pruning are likely to contribute to these sex-specific synaptic abnormalities, but are mindful of the possibility that other processes, such as changes in connectivity between the entorhinal cortex and the dorsal hippocampus, may also play a role.

### LB Impairs Microglial Mediated Synaptic Pruning at P17 but not P33 Adolescent mice

Consistent with previous findings (Dayananda et al., 2022), we determined that LB causes severe deficits in microglial mediated synaptic pruning during the second and third weeks of life, and clarified that these abnormalities are no longer present in P33 adolescent LB mice. Interestingly, the numbers of PSD95 puncta inside microglia or inside the phagosome were roughly two-fold greater in adolescent (P33) LB compared to CTL mice. We hypothesize that higher rates of synaptic pruning in adolescent LB mice may serve as a compensatory mechanism to minimize the impact of deficits earlier in life but may also lead to “over pruning” and cognitive deficits later in life. We show that the rate of synaptic engulfment is 7.6-fold higher at P17 compared to P33 in CTL mice. This finding is consistent with previous work showing that microglial mediated synaptic pruning peaks in the hippocampus during the second and third weeks of life and that perturbation of this process leads to reduced connectivity in the Schaffer collaterals later in life (Filipello et al., 2018; Scott-Hewitt et al., 2020; Zhan et al., 2014). Zhan et al. (2014) have suggested that synaptic pruning is necessary for the formation of multisynapse boutons 1(MSB1s). However, since MSB1s account for only 5% of all synaptic boutons, a reduction in this small population of synaptic connections is unlikely to fully account for the synaptic and cognitive deficits induced in response to abnormal microglial synaptic pruning in the developing hippocampus. We propose that spine formation and maturation is a competitive intracellular process for limited neuronal resources. The removal of non-functional or weak spines during the second and third weeks of life is necessary to support the formation of mature mushroom spines that are necessary for normal hippocampal function. The maturity index provides a highly sensitive structural “footprint” for detecting earlier deficits in microglial-mediated synaptic pruning. Importantly, both LB and transient ablation of microglia halve the synaptic maturity index, and this index is strongly correlated with contextual fear conditioning.

### Transient Elimination of Microglia Leads to Similar Sex-Specific Deficits Observed in LB Adolescent Mice

This study is the first to investigate the effects of transiently ablating microglia during the second and third weeks of life on hippocampal function in adolescent male and female mice. Our findings indicate that microglia play a more prominent role in modulating synaptic plasticity and contextual fear conditioning in male compared to female mice. These findings are supported by research indicating that genomic differences between male and female microglia become apparent during the second week of life (Hanamsagar et al., 2017; Thion et al., 2018). Additionally, it has been shown that microglia have sex-specific impact on pain sensitivity (Sorge et al., 2015), obesity (Dorfman et al., 2017), and neuroprotection against ischemic strokes (Villa et al., 2018). Our findings are also consistent with studies showing that transient perturbation of microglial function during a critical period of development results in long-term alterations in synaptic plasticity and behavior (Bolton et al., 2022; Favuzzi et al., 2021; Garvin and Bolton, 2022). For example, localized ablation of microglia in the prefrontal cortex of 6-week-old male adolescent mice resulted in impaired working memory, decreased density of mature mushroom spines, and reduced glutamatergic synapse density in adulthood. These abnormalities were not observed when microglia were ablated in adult animals (Schalbetter et al., 2022). Unfortunately, most ablation studies have been conducted exclusively in male mice. However, the few studies that have examined both sexes found significant differences between males and females (Bolton et al., 2022; Favuzzi et al., 2021; Garvin and Bolton, 2022). The similarities between outcomes in adolescent LB mice and adolescent DTA mice (Figs 1 & 3) support the idea that transient ablation of microglia contributes to sex differences in contextual fear conditioning, maturity index, glutamatergic synapses, and local functional connectivity in the hippocampus. The mechanisms responsible for the sex differences reported here in normally developing DTA mice are yet to be elucidated, but it is possible that male mice have higher rates of synapse formation compared to females. Assuming comparable rates of synapse elimination in males and females (Figure 2 and Figure S6), elimination of microglia would thus be expected to cause a more prominent reduction in the maturity index in males.

This is the first study to use rsfMRI to evaluate local functional connectivity in an animal model of ELA. This approach provides an unbiased method to identify brain regions with presumed differences in local connectivity (Tomasi and Volkow, 2010). However, to the best of our knowledge, this is first example in which reduced local functional connectivity is linked to reduced structural synaptic abnormalities. Using this approach, we identified several other brain regions that show sex-specific changes in local connectivity in adolescent LB mice. These include the entorhinal cortex, striatum, basolateral amygdala, nucleus accumbens, and the hypothalamus. Additional work is required to further characterize the impact of LB and sex on synaptic changes in these brain regions. It would be valuable to determine whether similar sex-specific changes can be detected in humans exposed to various forms of ELA. Finally, the observation that transient ablation of microglia reduces local functional connectivity in several brain regions of normally developing adolescent mice suggests that this approach may provide an indirect method to monitor microglial-mediated synaptic pruning during early development in humans.

### Chemogenetic Activation of Microglia During the Second and Third Weeks of Life Normalizes Microglial-Mediated Synaptic Pruning and Hippocampal Function

Chemogenetic activation of microglia from P13-P17 was able to normalize microglial phagocytic activity at P17 in LB male and female pups to levels seen in CTL. This, in turn, reversed the deficits in contextual freezing and synaptic abnormalities observed in adolescent LB males. These results align with recent findings indicating that chemogenetic activation of microglia during the first week of life can restore normal glutamatergic innervation of CRH-positive cells in the hypothalamus, adrenal size, and stress reactivity in male LB mice (Bolton et al., 2022). Our findings demonstrate that increasing levels of microglial-mediated synaptic pruning during the second and third weeks of life is sufficient to normalize the cognitive and synaptic abnormalities observed in adolescent LB males and suggest that interventions enhancing microglial-mediated synaptic pruning in childhood may offer an effective therapy for early adversities characterized by low functional connectivity.

### LB Increases MEGF10 and Astrocyte-Mediated Synaptic Pruning in the Developing Hippocampus of Female Mice

Previous work has shown that microglial mediated synaptic pruning in the hypothalamus is impaired in 10-days old LB male, but not LB female, pups (Bolton et al., 2022). This was not the case during peak synaptic pruning in the hippocampus where both male and female mice showed pronounced deficits in microglial phagocytic activity in vivo and ex vivo (Dayananda et al., 2022). Given that female adolescent mice show minimal synaptic and cognitive deficits and that astrocytes can compensate for deficits in microglial-mediated synaptic pruning (Chung et al., 2013; Damisah et al., 2020; Konishi et al., 2020; Lee et al., 2021; Perez-Catalan et al., 2021), we assessed the effects of LB on synaptic pruning in astrocytes located in the stratum radiatum. We found that LB increased the number of PSD95 puncta in astrocytes from female, but not male, mice, an increase that was associated with upregulation of the MEGF10 receptor. This is consistent with previous work showing that MEGF10 is necessary to support normal synaptic pruning in astrocytes (Chung et al., 2013; Lee et al., 2021) and suggest a mechanism by which female LB mice are able to compensate for microglial-mediated synaptic deficits. Future studies will test whether cell-specific deletion of MEGF10 in astrocytes render females more sensitive to LB-mediated changes in hippocampal function. Furthermore, clarifying the mechanisms by which females upregulate the MEGF10 receptor, may offer novel strategies for minimizing hippocampal-dependent synaptic and cognitive deficits in male LB mice.

### Conclusions

A better understanding of the underlying pathophysiology through which ELA alters hippocampal function in both males and females is essential for the development of more effective therapies. Here we show that LB causes more severe deficits in contextual freezing and synaptic plasticity in adolescent male compared to female mice. At P17, when synaptic pruning peaks in the hippocampus, both male and female pups exhibit a fourfold reduction in microglial-mediated synaptic pruning. This is no longer the case at P33 when the rate of microglial-mediated synaptic pruning is significantly lower. Transient ablation of microglia during the second and third weeks of life leads to sex-specific deficits similar to those observed in LB adolescent mice. Chemogenetic activation of microglia during the same period is sufficient to reverse the contextual and synaptic abnormalities observed in LB male adolescent mice. Finally, levels of MEGF10 and synaptic engulfment are elevated in 17-day-old LB females, suggesting a potential compensatory mechanism for reducing the microglial-mediated abnormalities in LB females. These studies highlight a novel role for glial cells in mediating sex-specific hippocampal deficits in a mouse model of ELA.

## AKNOWLEDGMENTS

This work was supported by: NIMH R01MH119164, NIMH R01MH118332, and the Clinical Neuroscience Division of the VA National Center for PTSD.

## Supplemental Information

**Fig S1.**
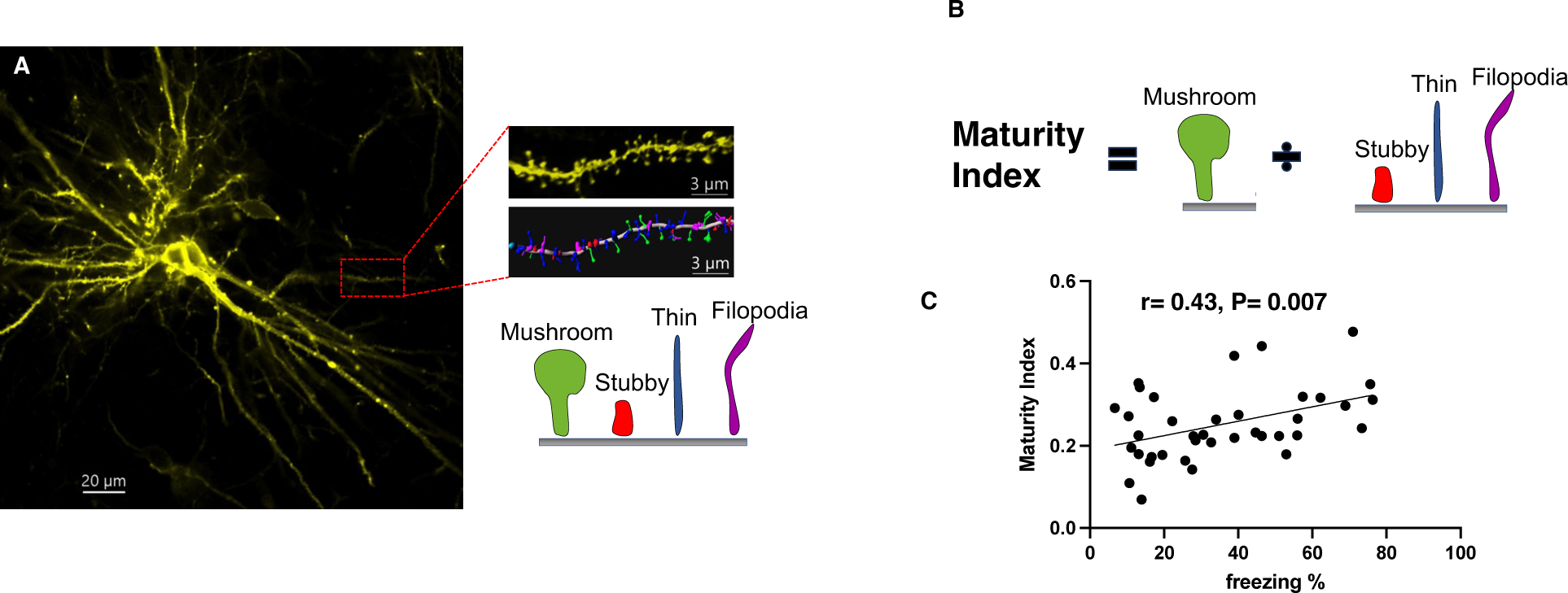
The Maturity Index Correlates with Contextual Fear Conditioning. (A) A representative low magnification image of CA1 pyramidal neuron labeled with DiI (Left). High magnification of a secondary apical dendrite and an Imaris color-coded model of spine morphology. (B) The ratio between the number of mushroom spines and all other spines was calculated for each apical dendrite and then averaged across 5-6 dendrites to quantify the maturity index for each mouse. (C) The maturity index is correlated with contextual fear conditioning in adolescent CTL and LB mice. Significant correlation between the maturity index and contextual fear conditioning was also seen in WT and DTA adolescent mice (r= 0.74, P= 0.0027, n= 14), and in adolescent mice in which microglia were chemogenetically activated from P13-17 (r= 0.87, P= 0.001, n= 16).

**Fig S2.**
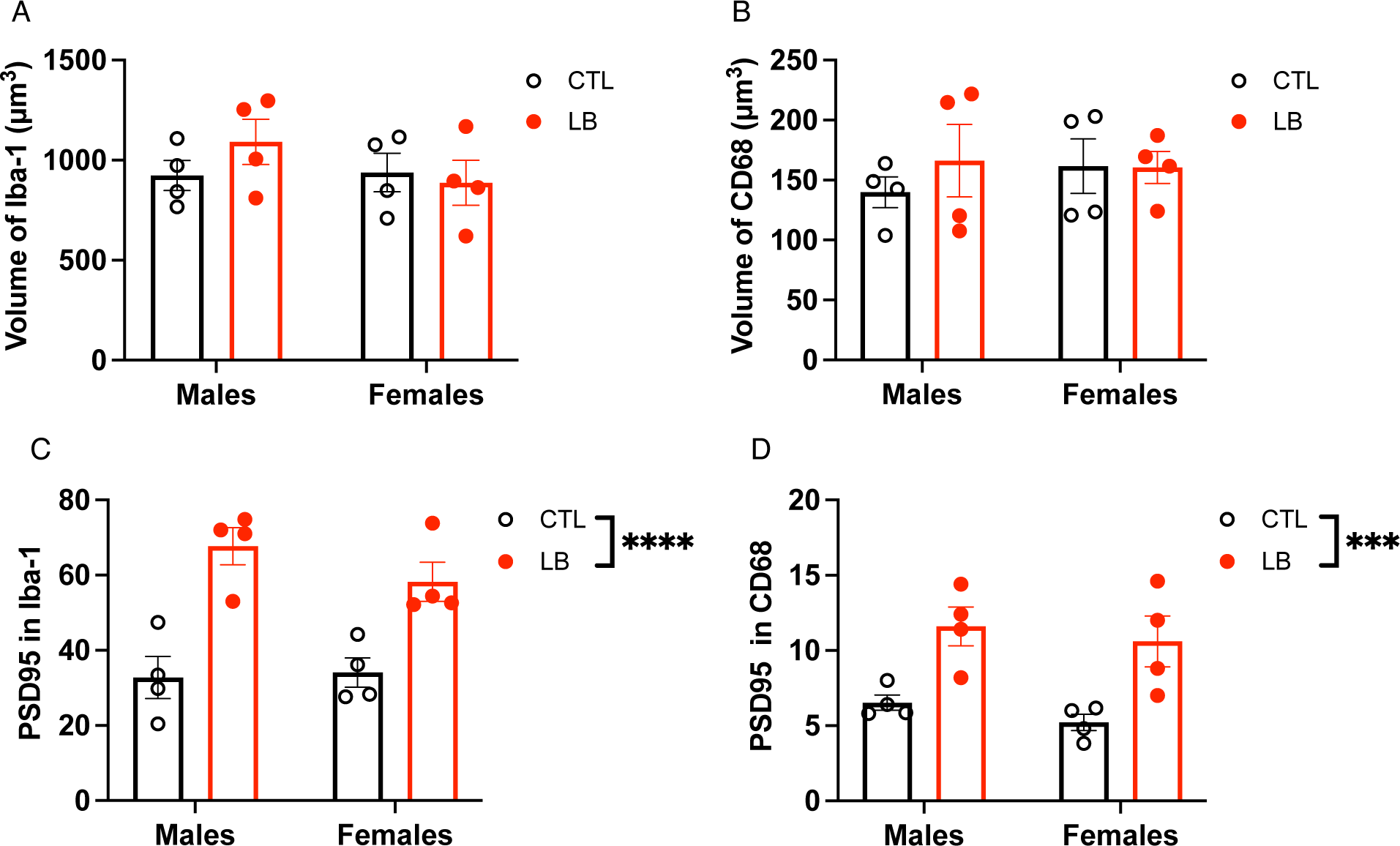
LB Increases Microglial Phagocytic Activity in Adolescent P33 mice. CTL and LB adolescent mice were perfused at P33 to assess microglial volume, CD68 volume, number of PSD95 puncta inside microglia, and number of PSD95 puncta inside CD68. For representative images see figure 2. (A) Microglial volume. Rearing: F (1, 12) = 0.344, P= 0.57, Sex: F (1, 12) = 0.89, P= 0.36, Interaction: F (1, 12) = 1.19, P= 0.29 (B) CD68 phagosome volume inside microglia. Rearing: F (1, 12) = 0.36, P= 0.56, Sex: F (1, 12) = 0.14, P= 0.71, Interaction: F (1, 12) = 0.42, P= 0.53. (C) PSD95 puncta inside microglia. Rearing: F (1, 12) = 35.54, P< 0.0001, Sex: F (1, 1. 12) = 0.67, P= 0.43, Interaction: F (1, 12) = 1.18, P= 0.30 (D). PSD95 puncta inside CD68. Rearing: F (1, 12) = 21.62, P= 0.0006, Sex: F (1, 12) = 1.054, P= 0.32, Interaction: F (1, 12) = 0.019, P= 0.89. Error bars represent mean ± SEM. Two-way ANOVA, ***p< 0.001, ****p< 0.0005.

**Fig S3.**
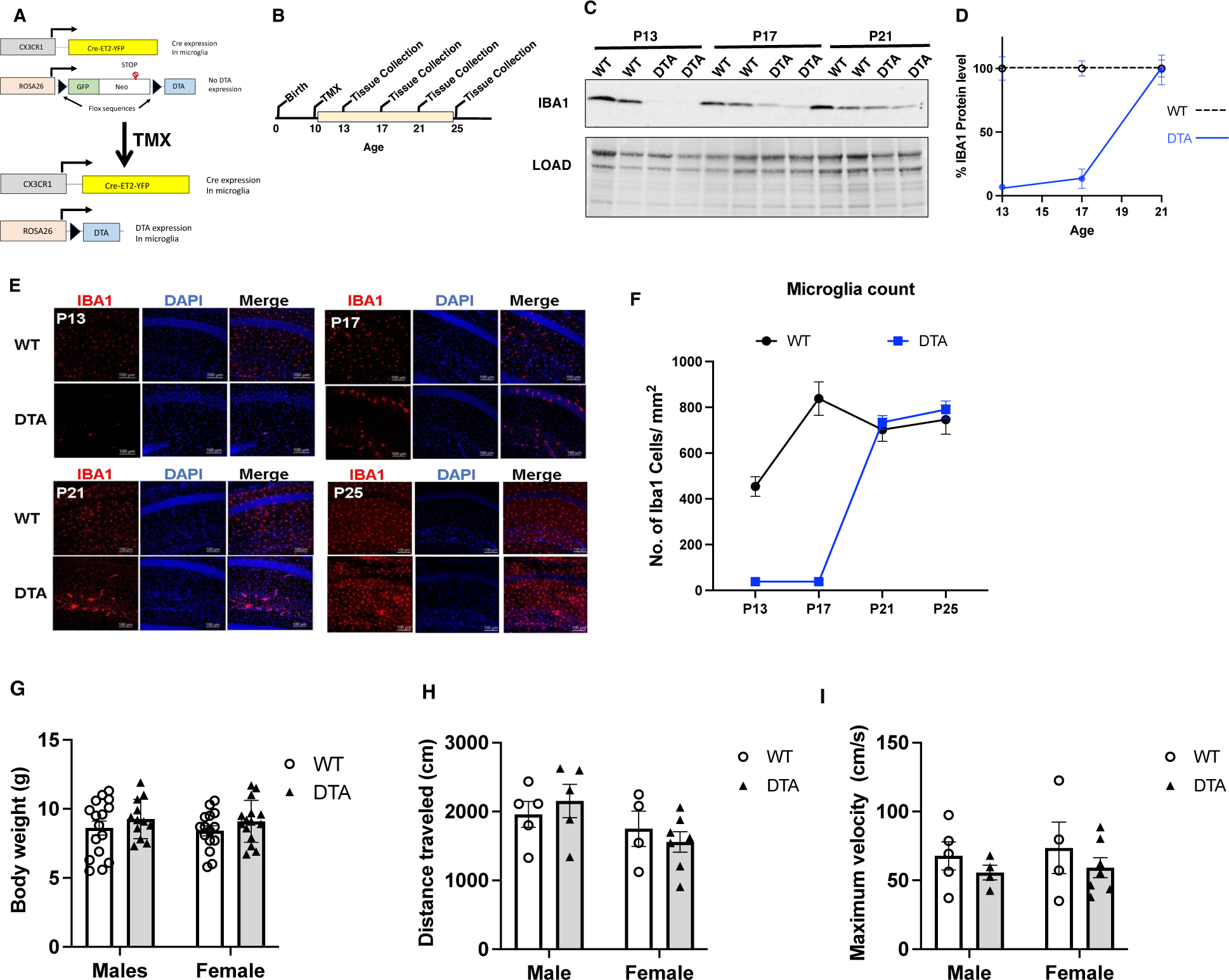
Transient Microglial Ablation During the Second and Third Weeks of Life Does not Impact Weight or Mobility at P17. (A) Schematics of ablation strategy. Tamoxifen (TMX) induces Cre mediated removal of a neomycin-GFP cassette containing a stop codon allowing for the expression of the diphtheria toxin A (DTA) subunit in microglia. (B) Experimental timeline. TMX (30mg/kg) was administered i.p. at P10 and tissue was collected at P13, P17, P21 and P26. (C-D) Western blot and quantification of Iba1 in the hippocampus (N= 5-7 mice per age and genotype, 50% females). (E-F) Confocal images and quantification of the number of Iba1-positive cells in the stratum radiatum (N= 6-8 mice per age and genotype, 50% females). Effects of genotype (WT vs DTA), sex, and their interaction in body weight (G), total distance moved (H) and maximum in the open field test velocity (I). Error bars represent mean ± SEM.

**Fig S4-.**
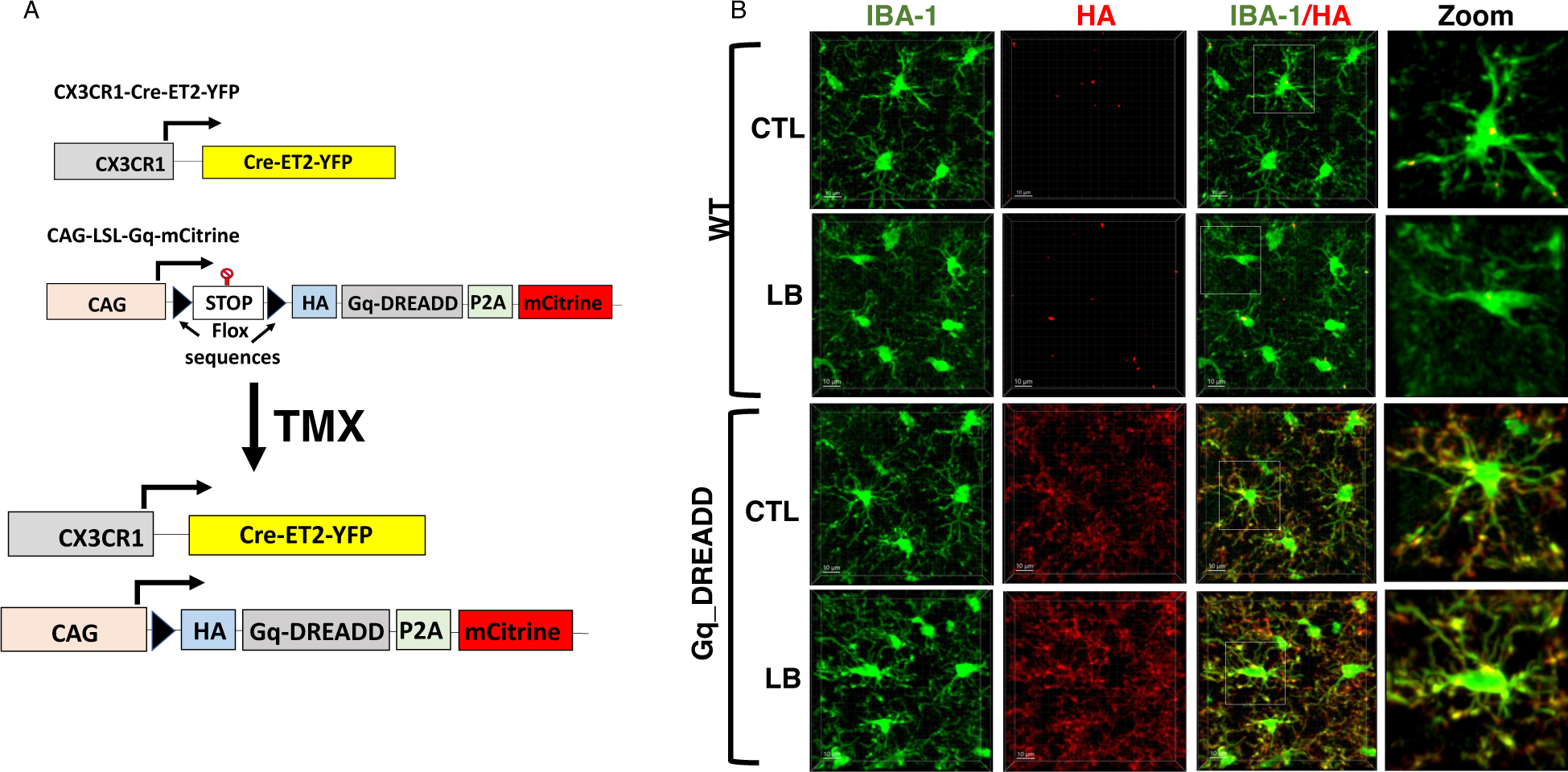
HA-Gq-DREADD expression in P17 Microglia. (A) Schematic illustration of the tamoxifen induced expression of HA-Gq-DREADD in CX3^Cre-ET2^-Gq-DREADD mice. (B) Gq and WT pups were administered tamoxifen i.p. at P10 (30mg/kg), followed by daily CNO injections P13-17 and perfused at P17. Tissue was then stained with anti-HA antibody (1:1000; BioLegend, Cat. #901501) and anti-Iba1 antibodies (1:500; Wako, Cat. #019-19741). HA-Gq-DREADD expression was detected in microglia from Gq but not WT littermates.

**Fig S5-.**
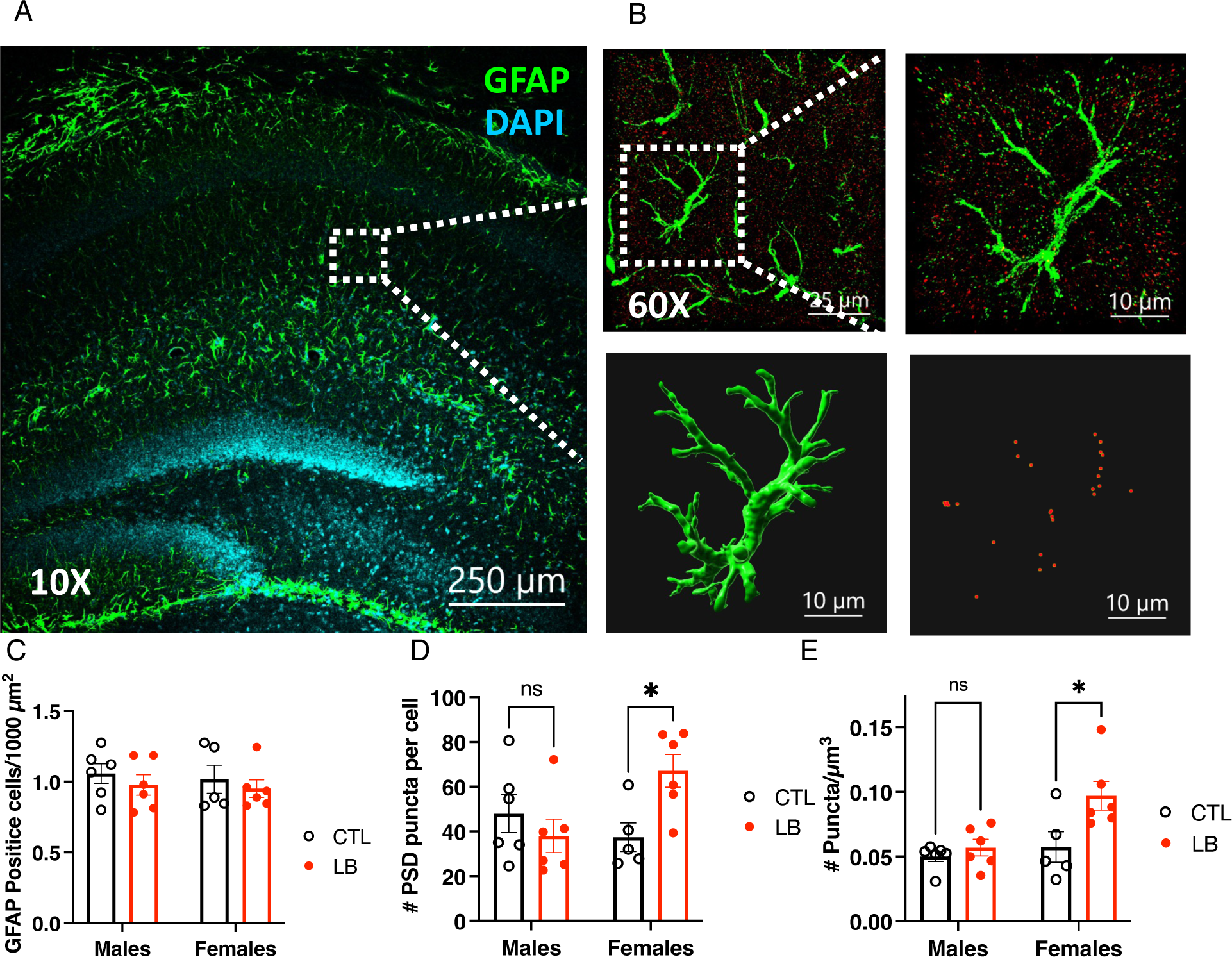
LB Increases Phagocytic Activity in Female Astrocytes Located in the Stratum Radiatum. (A) Low magnification of GFAP-positive cells in the dorsal hippocampus. (B) Higher magnification of GFAP (green) and PSD95 (red) staining and Imaris models of a GFAP-positive astrocyte located in the stratum radiatum and PSD95 puncta inside the cell. (C) Density of GFAP-positive cells in the stratum radiatum. Rearing: F (1, 19) = 0.95, P= 0.34, Sex: F (1, 19) = 0.19, P= 0.67, Interaction: F (1, 19) = 0.009, P= 0.92. (D) Number of PSD95 puncta per GFAP-positive cell, Rearing: F (1, 19) = 1.694 P= 0.21, Sex: F (1, 19) = 1.48, P= 0.24, Interaction: F (1, 19) = 6.82; P= 0.017, Males: P= 0.58, Females: P= 0.028. (E) Density of PSD95 puncta in GFAP-positive astrocytes, Rearing: F (1, 19) = 7.18, P= 0.014, Sex: F (1, 19) = 7.54; P= 0.013, Interaction: F (1, 19) = 3.6, P= 0.073. Males: P= 0.82, Females P= 0.010. Error bars represent mean ± SEM. *p< 0.05.

**Fig S6.**
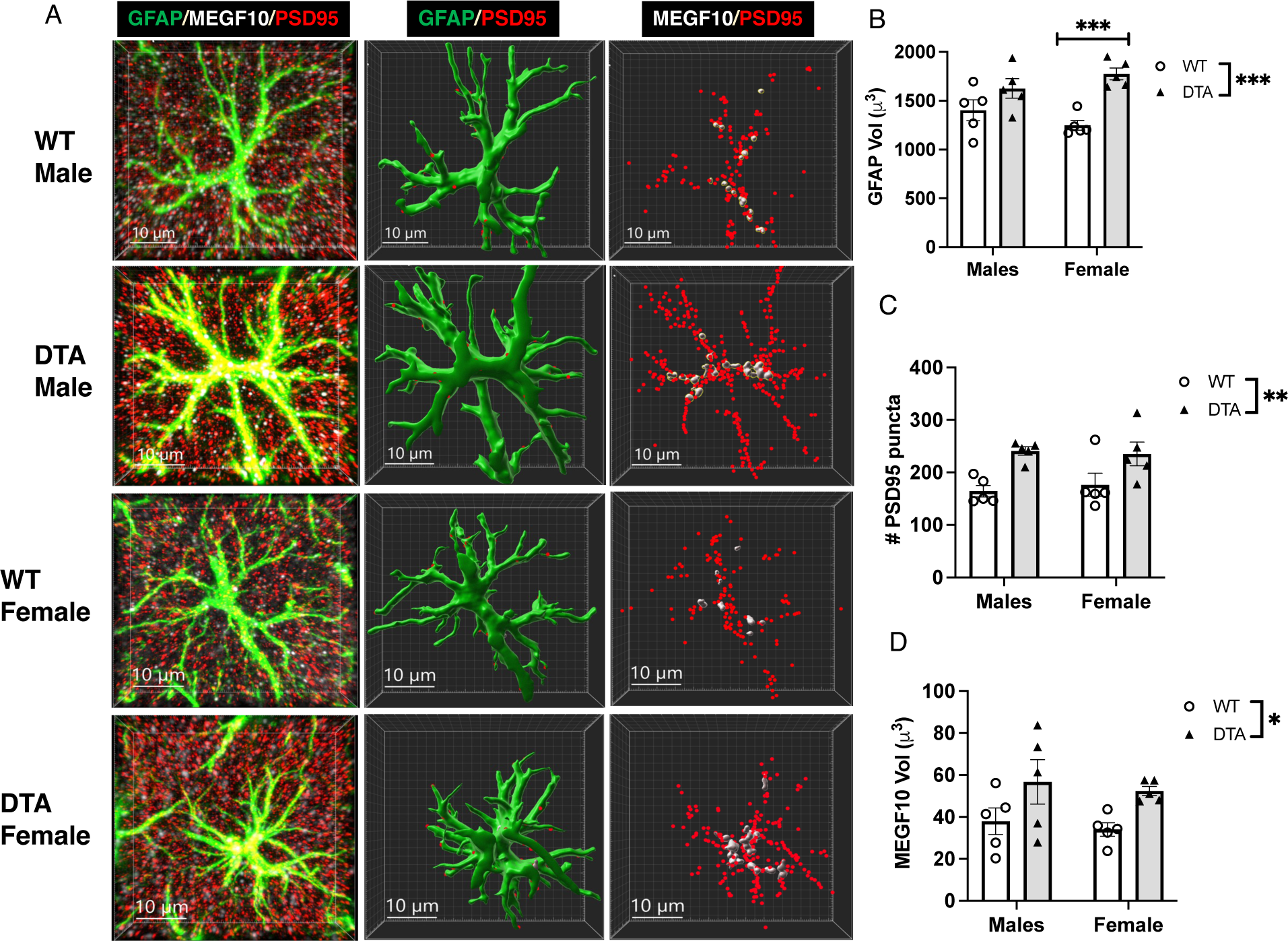
Microglial Ablation Increases PSD95 Puncta and MEGF10 Expression in Normally Developing Male and Female Pups. WT and DTA P10 pups were administered tamoxifen i.p. at P10 and processed at P17 to assess phagocytic activity and MEGF10 levels in GFAP-positive astrocytes located in the stratum radiatum. (A) Representative confocal images and Imaris models. (B) Cell volume, Rearing: F (1, 16) = 20.34, P=0.0004, Sex: F (1, 16) = 0.00037 P= 0.98, Interaction: F (1, 16) = 3.33, P=0.086, Sidak post-hoc, males: P= 0.15, females: P= 0.0008. (C) Number of PSD95 puncta inside GFAP-positive astrocytes, Rearing F (1, 16) = 15.59, P=0.0012, Sex: F (1, 16) = 0.035, P= 0.85, Interaction: F (1, 16) = 0.26, P= 0.62, Sidak post-hoc, males: P= 0.11, females: P= 0.12. (D) MEGF10 staining in astrocytes, Rearing: F (1, 16) = 8.2, P= 0.011, Sex: F (1, 16) = 0.41, P= 0.53, Interaction: F (1, 16) = 0.00075, P= 0.97, Sidak post-hoc, males: P= 0.012, females: P= 0.053. Error bars represent mean ± SEM. *p< 0.05, **p< 0.01, ***p< 0.001.

